# *Drosophila* hemocytes recognize lymph gland tumors of *mxc* mutants and activate the Toll pathway in reactive oxygen species-dependent manner

**DOI:** 10.1101/2021.12.17.473119

**Authors:** Suzuko Kinoshita, Kazuki Takarada, Yoshihiro H. Inoue

## Abstract

Mechanisms of cancer cell recognition and elimination by the innate immune system remains unclear. Circulating hemocytes are associated with the hematopoietic tumors in *Drosophila mxc*^*mbn1*^ mutant larvae. The innate immune signalling pathways are activated in the fat body to suppress the tumor growth by inducing antimicrobial peptides (AMP). Here, we investigated the regulatory mechanism underlying the activation in the mutant. Reactive oxygen species accumulated in the hemocytes due to induction of dual oxidase and its activator. The hemocytes were also localized on the fat body. These were essential for transmitting the information on tumors toward the fat body to induce AMP expression. Regarding to the tumor recognition, we found that matrix metalloproteinase 1 (MMP1) and MMP2 were highly expressed in the tumors. Ectopic expression of MMP2 was associated with AMP induction in the mutants. Furthermore, the basement membrane components in the tumors were reduced and ultimately lost. The hemocytes may recognize the disassembly in the tumors. Our findings highlight the underlying mechanism via which macrophage-like hemocytes recognize tumor cells and relay the information toward the fat body to induce AMPs. and contribute to uncover the immune system’s roles against cancer.

**SUMMARY STATEMENT:** *Drosophila* blood cells transmit information regarding the existence of tumor cells to the fat body and induce the production of anti-tumor peptides via the generation of reactive oxygen species.

## INTRODUCTION

Insects rely fundamentally on the innate immune system to escape parasite-microbial infection, as they lack an acquired immune system (Brennan and Anderson, 2004; Buchon et al., 2014; Hoffmann, 2003). The innate immunity of *Drosophila* is roughly divided into two types, humoral and cell-mediated immune responses. Humoral immunity involves the use of anti-microbial peptides (AMPs) which are synthesized in the fat body and secreted into the hemolymph (Hoffmann and Reichhart, 2002; Tzou et al., 2002). Seven distinct AMPs, together with their isoforms, have been identified in *Drosophila melanogaster* (Lemaitre and Hoffmann, 2007; Martinelli and Reichhart, 2005). *Drosophila* AMPs can not only destroy invading microorganisms but can also suppress the progression of several types of tumors in the larvae (Araki et al., 2019, Bilder et al., 2021, Parvy et al., 2019). Araki and colleagues have reported that overexpression of any one of the five AMPs studied enhanced apoptosis in mutant lymph glands (LGs), whereas no apoptotic signals were detected in the controls. Expression of these AMP genes is regulated by two major innate immune signalling pathways (Buchon et al., 2014; Hoffmann and Reichhart, 2002; Tzou et al., 2002). The first one is the Toll-mediated pathway, which is activated mainly by gram-positive bacteria and fungi (Lemaitre et al., 1997). Infection of these microbes initiates activation of consecutive serine protease cascades, which ultimately lead to the production of an active ligand called Spätzel (Jang et al., 2006; Valanne et al., 2011). The active ligand binds and activates the transmembrane receptor, Toll, (Buchon et al., 2014; Morisato and Anderson, 1994). This results in the assembly of several cytoplasmic proteins, including Pelle, and stimulates phosphorylation of Cactus, the negative regulator of the NF-κB family of transcription factors Dorsal (Dl) and Dorsal-related immunity factor (Dif) (Sun et al., 2002). In the absence of infection, the inhibitor prevents Dl from entering the nucleus. Once Cactus is degraded in response to infection, these transcription factors are free to translocate into the nucleus and induce the expression of genes encoding antimicrobial peptides such as Drosomycin (Drs) (Fehlbaum et al., 1994; Hoffmann, 2003). The second innate immune signalling pathway is activated by infection of Gram-negative bacteria. These bacteria are recognised by peptidoglycan recognition protein (PGRP)-LC (Choe et al., 2002; Gottar et al., 2002), which leads to the activation of the signalling pathway mediated by a multiprotein complex containing Imd, FADD, and Dredd (Choe et al., 2005; Georgel et al., 2001; Lemaitre et al.,1995). Subsequently, the protein complex phosphorylates the Relish transcription factor (Rel), which triggers the cleavage of Rel (Kleino et al., 2005; Silverman et al., 2000). The resultant N terminal Rel drives the expression of several antimicrobial peptides, such as Diptericin (Dpt) (Stöven et al., 2000; Stöven et al., 2003).

Cellular immune response also plays a significant role in protecting against invading pathogens in *Drosophila*. In case of infection, phagocytosis by macrophage-like hemocytes and encapsulation and melanisation by other types of hemocytes come into play. The circulating hemocytes arise from two distinct haematopoietic tissues, the embryonic head mesoderm and a specialised organ called the LG at a later larval stage (Lanot et al., 2001; Araki et al., 2019). The LG contains haematopoietic progenitor cells called pro-hemocytes, which can give rise to three types of hemocytes: plasmatocytes, lamellocytes, and crystal cells (Evans et al., 2003; Govind, 2008). *Drosophila* can respond to invading pathogenic microorganisms, as well as to tumor cells in the body (Wang et al., 2014). The hemocytes associate with the tumors and activate the innate immune system, thereby restricting tumor growth (Parisi et al., 2014; Pastor-Pareja et al., 2008; Wang et al., 2014). A series of studies have investigated the crosstalk between cellular immune response and tumors in *Drosophila* (Arefin et al., 2017; Kalamarz et al., 2012; Kim and Choe, 2014). Two AMPs, Drosomycin and Defensin, were incorporated by circulating hemocytes associated with the LG tumors (Araki et al., 2019). These results suggested that AMPs possess a specific cytotoxic property, because of which they induce apoptosis exclusively in the tumors. Another subsequent study reported that Defensin can suppress tumor progression by targeting phosphatidyl serine on tumor cell membranes and inducing cell death (Parvy et al., 2019). However, the mechanism via which the innate immune system detects tumor cells still remain unknown. Furthermore, the mechanism via which the information is transmitted from the LG toward the fat body to induce AMPs in the tissue is not understood.

*Drosophila* harbouring mutation in an allele of *multi sex combs* (*mxc*), *mxc*^*mxc*^, has been considered a haematopoietic tumor model (Remillieux-Leschelle et al., 2002; Shrestha and Gateff, 1982; Araki et al., 2019, Kurihara et al., 2020, Kurihara et al., 2020b). The *mxc*^*mbn1*^ mutant, which harbours a loss of function mutation in *mxc*, exhibits hyperplasia in the LG. LG cells isolated from the mutant larvae proliferated further, invaded host tissues, and ultimately killed the host when implanted into normal adult abdomens (Remillieux-Leschelle et al., 2002; Kurihara et al., 2020). Because of this, the mutant LG cells are considered malignant tumors (Kurihara et al., 2020; Remillieux-Leschelle et al., 2002; Shrestha and Gateff, 1982).

In this study, we aimed to determine the mechanism via which the LG tumors were recognized by the innate immune system and consequently activated to induce the expression of AMPs in the fat body of the *mxc*^*mbn1*^ larvae. We attempted to identify the factors that recognize the LG tumor cells in the larvae. Many studies have shown that altered proteolysis of the extracellular matrix (ECM) is related to unregulated tumor growth, tissue invasion, and metastasis. The matrix metalloproteinases (MMPs) are the most prominent proteinases associated with tumorigenesis (Kessenbrock, et al., 2010; Gobin et al., 2019). A previous study has also shown that the expression of the MMP1 proteinase, which cleaves the ECM proteins, was up-regulated in the *mxc*^*mbn1*^ mutant larvae, and that the proteinase was required for LG hyperplasia. Hence, the authors speculated that reduction in ECM due to MMP1 up-regulation influenced the tumor phenotype in the *mxc*^*mbn1*^ larvae (Kurihara et al., 2020). Here, we examined whether the MMPs were up-regulated in the mutant larvae. Next, we addressed the mechanism via which the information regarding the existence of tumor cells is transmitted to the fat body positioned separately from the LGs via the hemolymph. The findings in this study will improve our understanding regarding the mechanism via which the innate immune system in the fat body was activated apart from that in the tumors, and consequently, how the AMPs were secreted from the immune tissues.

## RESULTS

### Inhibition of the hyper-accumulation of ROS by antioxidant feeding reduced AMP induction and enhancement of LG hyperplasia in *mxc*^*mbn1*^ larvae

A previous study has shown that ROS were induced when a parasitic wasp injected its egg into a *Drosophila* larva, resulting in activation of Toll-mediated signalling in the larva and hemocyte-mediated killing of the egg (Louradour et al., 2017). Hence, we first investigated whether ROS had accumulated in *mxc*^*mbn1*^ larvae harbouring the LG tumor. We performed dihydroethidium (DHE) staining of hemocytes obtained from the hemolymph of *mxc*^*mbn1*^ larvae (*mxc*^*mbn1*^*/Y; He>GFPnls*) and of the cells from control larvae (*w/Y; He>GFPnls*) (Fig. 1A-C). We measured the DHE fluorescence in cells showing hemocyte-specific GFP fluorescence (Fig. 1A‴, B‴). DHE fluorescence increased significantly in *mxc*^*mbn1*^ hemocytes (median intensity 43116, n = 247) (*p* < 0.0001) compared to that in controls (median intensity value 21100, n = 357) (Fig. 1C). As the DHE fluorescence varied in intensity even among control hemocytes, we classified hemocytes into four classes [background level (Class I), weak (Class II), middle (Class III), and strong (Class IV)] according to the level of DHE fluorescence (Fig. 1D, E). Among the hemocytes in the hemolymph of control larvae (n = 247) 10.6% were in Class I, 38.1% in Class II, 25.2% in Class III, and 26.1% in Class IV. In contrast, 0.0 % of the *mxc*^*mbn1*^ hemocytes (n = 357) were in Class I, 12.6% in Class II, 30.4% in Class III, and 57.0% in Class IV. These results indicated that ROS had accumulated in the hemocytes of *mxc*^*mbn1*^ larvae. To confirm these results, we further investigated the fluorescence of the *gstD1-GFP* reporter, expression of which depends on ROS accumulation (Fig. S1). As expected, we observed significantly higher *gstD1* expression in the *mxc*^*mbn1*^ hemocytes than in the controls (Fig. S1 A’-D’, E), and the proportion of hemocytes showing stronger GFP fluorescence was higher in *mxc*^*mbn1*^ than that in the controls (Fig. S1 F, G). Next, we examined whether ROS also accumulated in *mxc*^*mbn1*^ larval LGs where hemocyte precursors were produced. Using the *gstD1-GFP* reporter, we compared the ROS levels in the LGs (Fig. S2A-C). Remarkably higher ROS levels were not observed in the mutant LGs (only 40% increase), compared to that in the controls (Fig. S2A’, S2B’, and S2C).

**Fig. 1.**
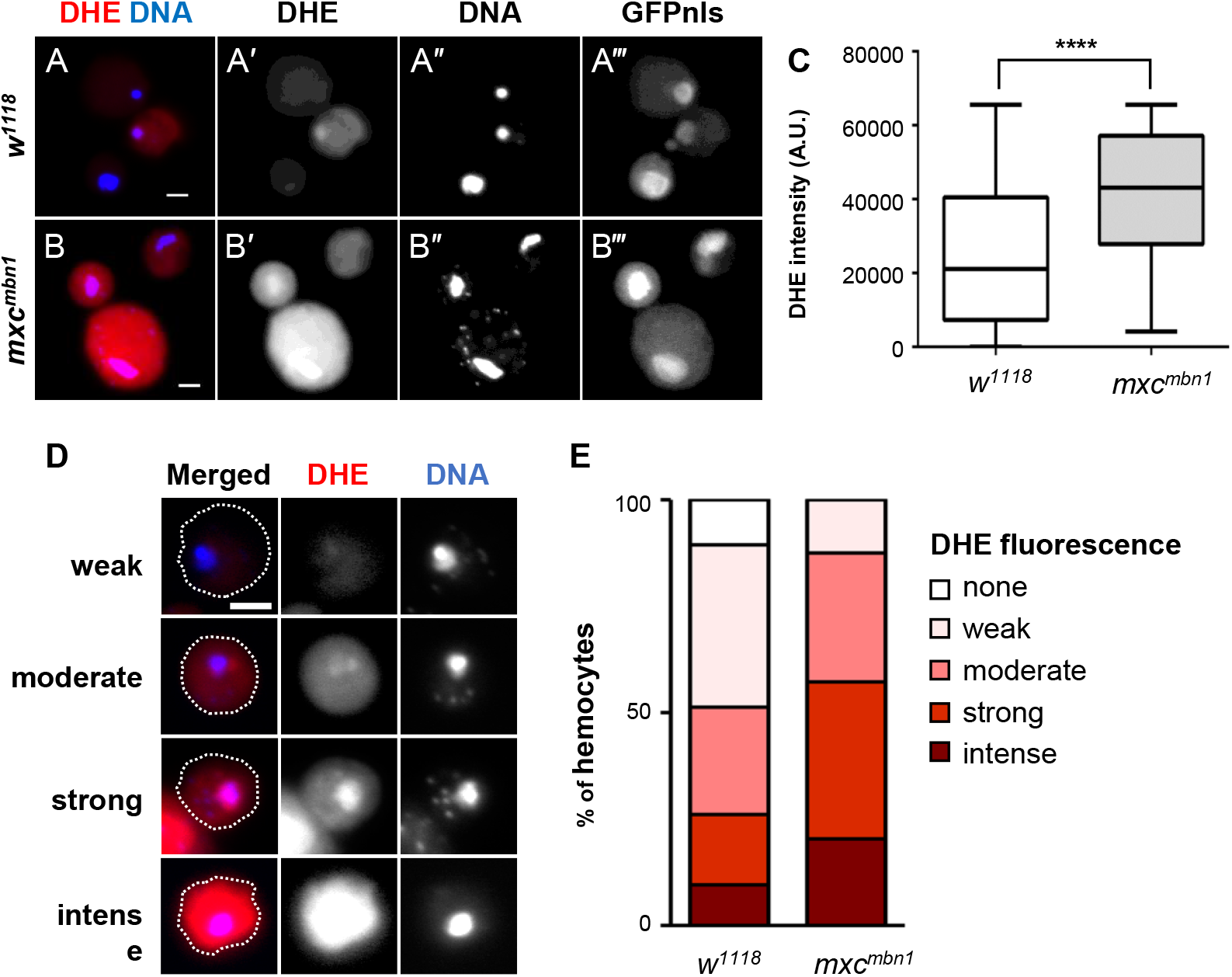
Hyper-accumulation of ROS in circulating hemocytes of *mxc*^*mbn1*^ mutant larvae. (A, B) DHE staining of circulating hemocytes from matured third instar larva (A; normal control (*w/Y; He>GFPnls*), B; *mxc*^*mbn1*^ mutant (*mxc*^*mbn1*^*/Y; He>GFPnls*). DHE fluorescence and DNA staining by DAPI are coloured in red (A, B, white in A’, B’) and in blue (A, B, white in A”, B”). A circulating hemocyte-specific GFP fluorescence (A‴, B‴). Bar: 5 μm. (C) Average arbitrary units of DHE fluorescence in circulating hemocytes (n≥247) from normal control (white bar) and *mxc*^*mbn1*^ mutant larvae (gray bar), respectively. Error bars represent standard error of mean. (\*\*\*\**p*<0.0001, Welch’s *t* test). (D) Typical images of the cells classified into the four classes according to the DHE fluorescence intensity (weak, moderate, strong, intense classes). The cell margins are encircled by dotted lines. Bar: 5 μm. (E) A percentage of each class of the DHE-stained circulating hemocytes from control (*w*) and *mxc*^*mbn1*^ larvae.

To understand the significance of ROS accumulation, we next investigated whether inhibition of the accumulation influenced the innate immune pathway and LG hyperplasia. Feeding on *Drosophila* diet containing an antioxidant, N-acetyl cysteine (NAC), eliminated the ROS in the larvae. We administered the NAC-supplemented diet to control larvae for 6 days until pupariation, and to *mxc*^*mbn1*^ larvae for 11 days. The level of *gstD-GFP* expression in hemocytes of control larvae (median intensity 33.46, n = 466) did not change distinctly, compared to that in the larvae not fed NAC (median intensity 30.46, n = 372) (*p* < 0.001, Fig. S1A, B, E). In contrast, the fluorescence in the mutant hemocytes decreased significantly after NAC feeding (median 33.49, n = 259) compared to that in the mutant larvae without NAC feeding (median 70.54, n = 174) (*p* < 0.0001, Fig. S1C, D, E). Similarly, we classified the hemocytes after NAC feeding into the four classes on the basis of GFP fluorescence (Fig. S1F, G). In control larvae without NAC feeding, 48.7% hemocytes were classified into Class I, 49.4% in Class II, and 1.9% in Class III and Class IV each. After NAC feeding, the percentages of control larvae hemocytes in Class I, Class II, and Class III shifted to 38.2%, 58.2%, and 3.6%, respectively (Fig. S1G). In *mxc*^*mbn1*^ larva without NAC feeding, we scored 16.7% cells in Class I, 33.3% in Class II, and 50% in Class III. Consistent with this, 47.5% hemocytes in the *mxc*^*mbn1*^ larvae were classified in Class I, 27.4% in Class II, and 25.1% in Class III after the feeding. In summary, NAC feeding suppressed ROS accumulation in hemocytes of *mxc*^*mbn1*^, but not in control larvae (Fig. S1G).

Next, we investigated whether the reduction in ROS level due to NAC feeding influenced activation of innate signalling in the fat body. The mRNA levels of *Drs* and *Def* in control larvae fed the NAC-containing diet decreased by 68.7% and 48.9% compared to the levels in non-fed larvae, respectively, although the mRNA level of *Dpt* increased by 50%. In contrast, in *mxc*^*mbn1*^ larvae, the average mRNA levels of all three genes in the fat body decreased by 81.5%, 38.3%, and 21.7%, respectively (Fig. 2A). Activation of the innate immune pathway in *mxc*^*mbn1*^ larvae was inhibited with reduction in ROS levels. Moreover, we investigated whether reduction in ROS levels affected LG hyperplasia in *mxc*^*mbn1*^ larvae (Fig. 2B-F). LG size did not differ significantly between control larvae fed NAC diet (median 0.04 mm^2^, n = 21) and those fed diet without NAC (median 0.04 mm^2^, n = 29) (Fig. 2F). However, the LG hyperplasia in *mxc*^*mbn1*^ larvae (median 0.53 mm^2^, n = 19) was significantly higher than that in the mutant larvae not fed the NAC diet (median 0.25 mm^2^, n = 28) (*p* < 0.0001) (Fig. 2F). We concluded that ROS accumulation in *mxc*^*mbn1*^ larvae was required for the activation of the innate signalling pathway in the fat body, resulting in suppression of LG hyperplasia.

**Fig. 2.**
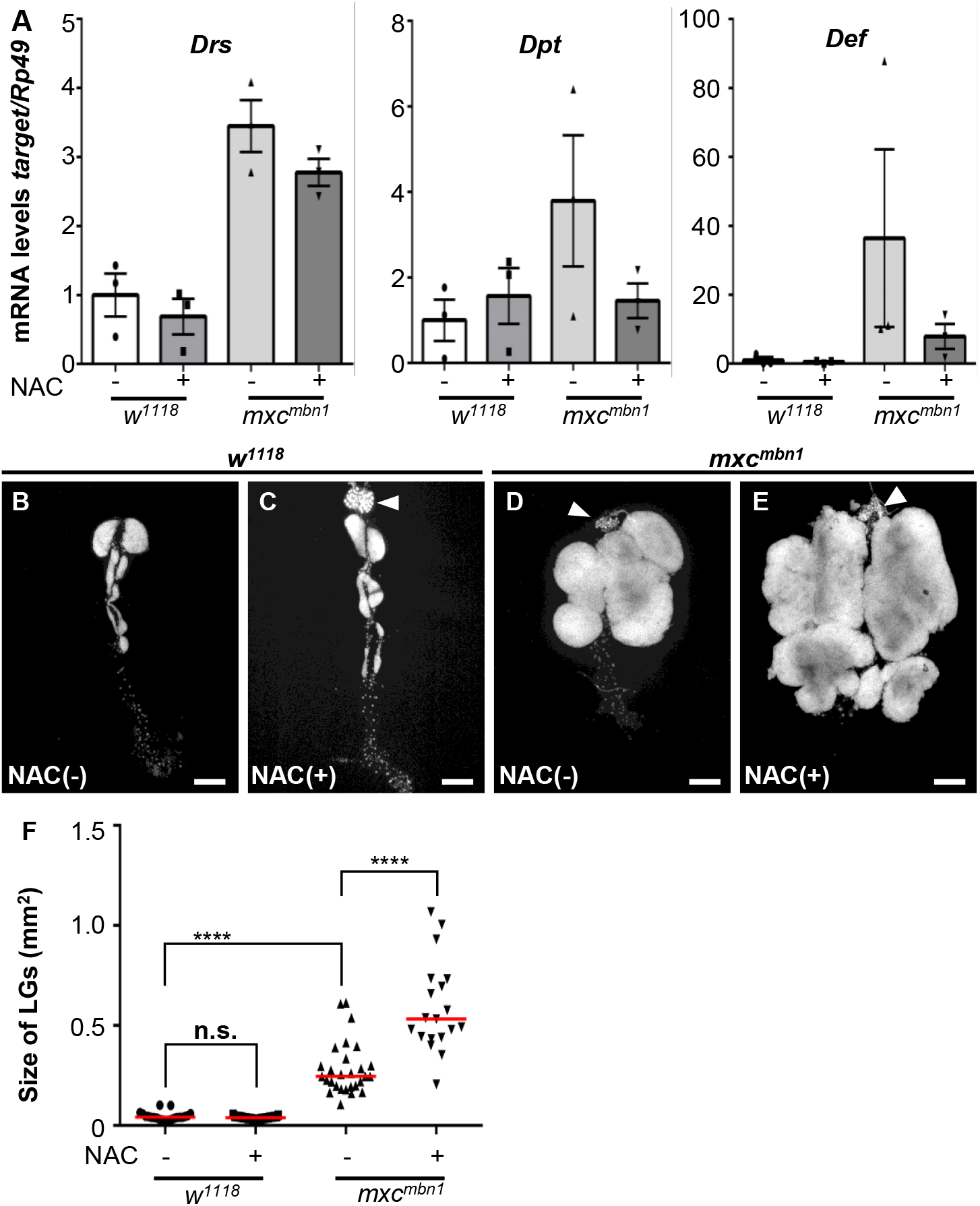
Reduced levels of AMPs and enlarged LG tumors in *mxc*^*mbn1*^ larvae fed on NAC. (A) The mRNA levels of *Drs, Def*, and *Dpt* in fat bodies quantified by qRT-PCR. Total RNA was prepared from control male larvae (*w*^*1118*^*/Y*) fed without NAC, with NAC, the mutant male larvae (*mxc*^*mbn1*^*/Y*) fed without, or with NAC. Average levels of the genes were calculated. (B-E) The DAPI-stained LGs from matured 3^rd^ instar larvae of control (*w*^*1118*^*/Y*) fed without NAC (B), with NAC (C), the mutant larvae (*mxc*^*mbn1*^*/Y*) fed without NAC (D), with NAC (E). Arrowhead indicates the ring gland linked to the LG. Bar; 200 µm. (F) Average size of the LGs from 3^rd^ instar larvae at mature stage (n≥19). The red lines indicate median size of the LG size. For statistical analysis of the differences between with or without NAC feeding, Kruskal-Wallis test followed by the Mann-Whitney U test using Bonferroni correction was performed (\*\*\*\**p*<0.0001, n.s.: not significant).

### *Duox* was up-regulated in *mxc*^*mbn1*^ larvae, and its depletion in hemocytes reduced the mRNA levels of AMP genes and increased LG hyperplasia

To identify the genes involved in ROS accumulation, we investigated whether the mRNA level of a unique gene encoding Dual Oxidase (*Duox*) changed in the mutant larvae. We quantified the mRNA level of *Duox* in the whole larvae, hemocytes in the hemolymph, fat body, and gut of control (*w*^*1118*^*/Y*) and *mxc*^*mbn1*^ larvae at the 3^rd^ instar stage (Fig. 3A-D). The average *Duox* mRNA level in whole *mxc*^*mbn1*^ larvae was 58.2%, while that in fat body was 54.8% of that of the control (Fig. 3A, C). In contrast, the level in *mxc*^*mbn1*^ larvae hemocytes was two-fold higher than that in the control (Fig. 3B); the level in gut of *mxc*^*mbn1*^ larvae hemocytes was also 50% higher than that in the control (Fig. 3D). Thus, *Duox* expression was stimulated in *mxc*^*mbn1*^ larvae, particularly in hemocytes and gut.

**Fig. 3.**
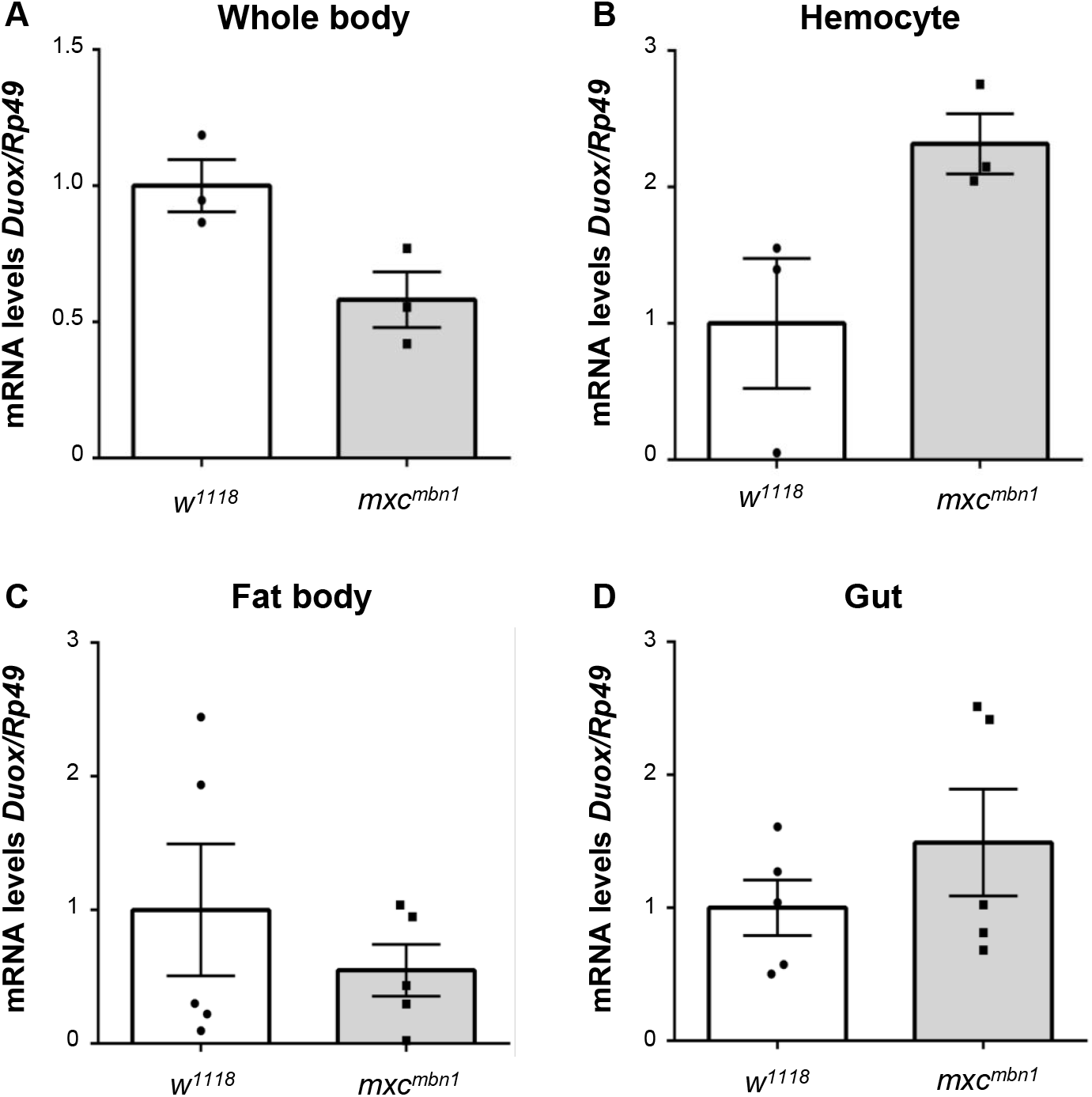
Increased levels of the *Duox* mRNA in circulating hemocytes in hemolymph of *mxc*^*mbn1*^. (A-D) Quantification of *Duox* mRNA by qRT-PCR using total RNAs prepared from whole 3^rd^ instar larvae at mature stage (A, n = 3), from hemocytes of the 3^rd^ instar larvae (B, n = 3), from fat bodies (C, n = 5), gut (D, n=5) as templates. A relative mRNA levels of *Duox* to an internal control (*Rp49*) was calculated. The average level in normal control (*w*^*1118*^) is presented as 1.0. The *Duox* mRNA levels are represented in the order of the control (white), and *mxc*^*mbn1*^ (gray).

Next, we investigated whether the up-regulation of *Duox* in *mxc*^*mbn1*^ hemocytes was required for activation of innate signalling in the fat body, and thereby suppression of LG hyperplasia. As *Duox* was efficiently depleted using dsRNA against its mRNA (Fig. S3), we performed qRT-PCR to quantify the mRNA levels of *Drs* and *Dpt* using total RNAs prepared from the fat body of the mutant larvae with or without hemocyte-specific depletion of *Duox* (Fig. 4A, B). In *mxc*^*mbn1*^ larvae harbouring hemocyte-specific *Duox* depletion, the mRNA levels of *Drs* and *Dpt* decreased by 35.2% and 13.3% compared to the levels in the mutant larvae without the depletion, respectively (Fig. 4A, B). These results indicated that *Duox* expression was required for the activation of the innate signalling pathways in *mxc*^*mbn1*^ larvae.

**Fig. 4.**
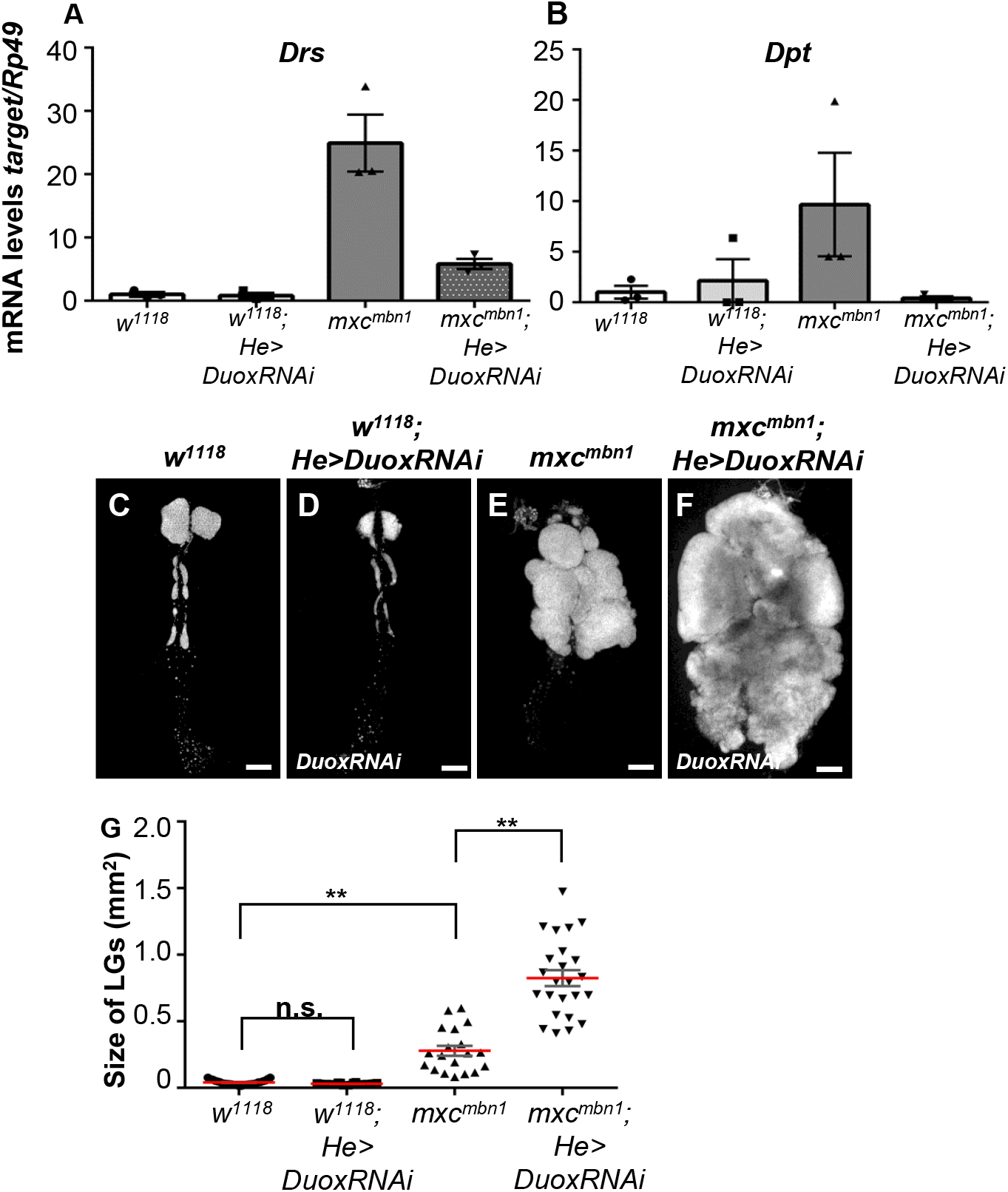
Reduced mRNA levels of two AMP genes and increased size of the LGs in 3^rd^ instar larvae harbouring hemocyte-specific depletion of the *Duox*. (A, B) Average mRNA levels of *Drs* (A) and *Dpt* (B) in fat bodies of 3^rd^ instar larvae at mature stage by qRT-PCR. Total RNA prepared from fat bodies of normal control larvae (*w*^*1118*^*/Y*), *mxc*^*mbn1*^ larvae (*mxc*^*mbn1*^*/Y*), *mxc*^*mbn1*^ larvae harbouring hemocyte-specific expression of *Duox* (*mxc*^*mbn1*^*/Y; He>DuoxRNAi*) was used as templates. Error bars represent standard error of mean. (C-F) The DAPI-stained LGs from matured 3^rd^ instar larvae of control (*w*^*1118*^*/Y*) (C), control larvae harbouring hemocyte-specific depletion of *Duox* (*w*^*1118*^ */Y; He>DuoxRNAi*) (D), *mxc*^*mbn1*^ larvae (*mxc*^*mbn1*^*/Y*) (E), *mxc*^*mbn1*^ larvae harbouring hemocyte-specific depletion of *Duox* (*mxc*^*mbn1*^*/Y; He>DuoxRNAi*)(F). Bar; 200 µm. (G) Average size of the LGs (n≥19). For statistical analysis of the differences in the LG size, one-way ANOVA with Scheffe’s multiple comparison test was performed (\*\**p*<0.01, n.s.: not significant). Red lines: average value. Error bars: standard error of mean.

Hence, we next investigated whether *Duox* depletion resulted in enhancement of the LG hyperplasia in the *mxc*^*mbn1*^ larvae. In *mxc*^*mbn1*^ larvae harbouring hemocyte-specific depletion of *Duox* (*mxc*^*mbn1*^*/Y; He>DuoxRNAi*), the average LG size (0.82 mm^2^, n = 24) was significantly larger than that of *mxc*^*mbn1*^ without the depletion (0.28 mm^2^, n = 19) (*p* < 0.01) (Fig. 4C-G). Collectively, *Duox* expression in hemocytes was required for activation of the innate signalling pathway and suppression of LG hyperplasia in *mxc*^*mbn1*^ larvae.

Furthermore, we investigated whether LG hyperplasia was suppressed by ectopic expression of *Duox* in the hemocytes (Fig. 5A-D). The average LG size (0.19 mm^2^, n = 21) in *mxc*^*mbn1*^ larvae with hemocyte-specific expression of Duox (*mxc*^*mbn1*^*/Y; He>Duox*) was significantly smaller than that in *mxc*^*mbn1*^ expressing GFP (*mxc*^*mbn1*^*/Y; He>GFP*) (0.39 mm^2^, n = 25) (*p* < 0.01) (Fig. 5E), indicating that ectopic expression of the ROS-producing enzyme resulted in suppression of LG hyperplasia in *mxc*^*mbn1*^ larvae. In summary, *Duox* expressed in circulating hemocytes played an essential role in suppression of LG hyperplasia.

**Fig. 5.**
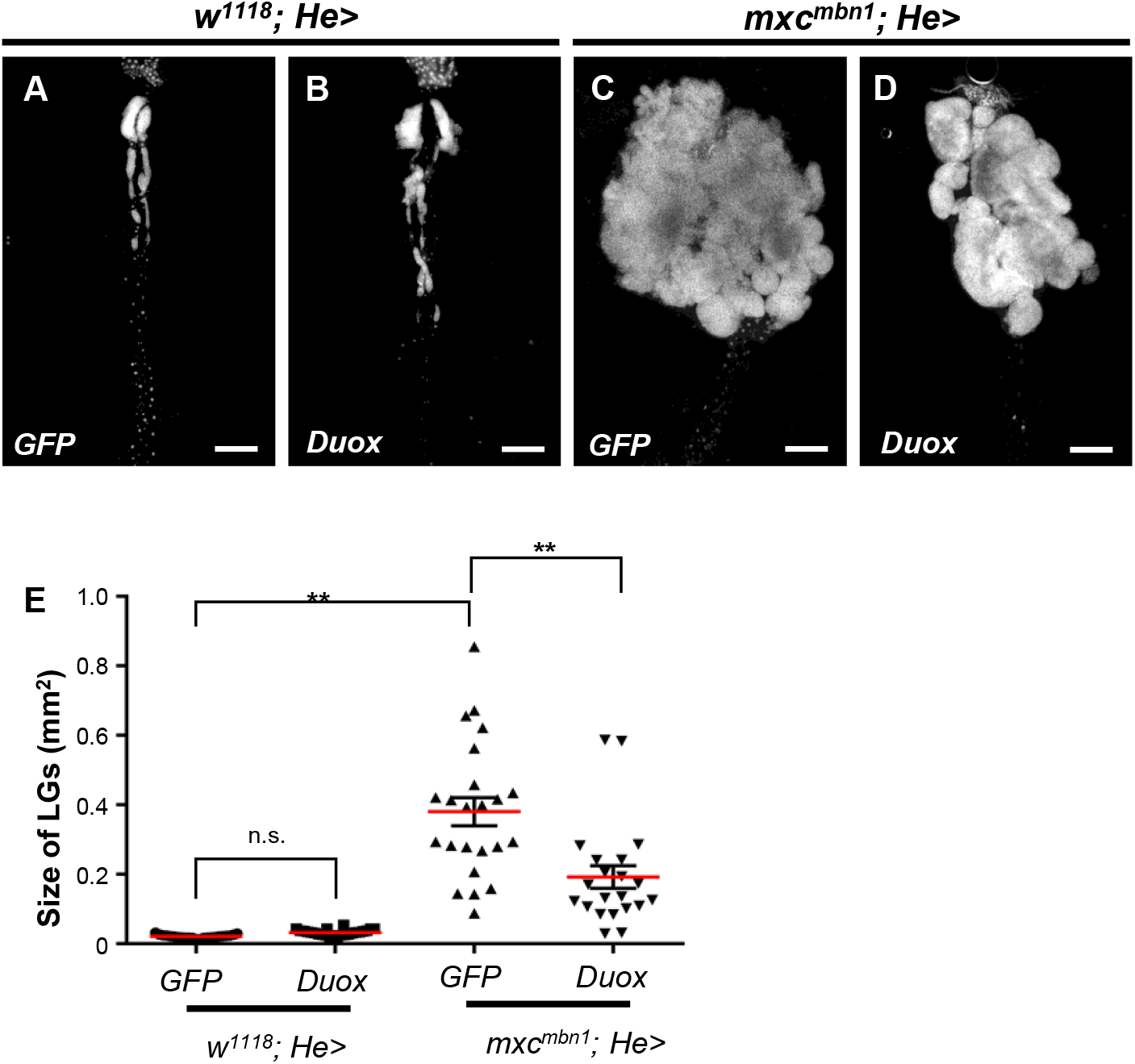
Reduced size of the tumorous LGs in in *mxc*^*mbn1*^ larvae harbouring hemocyte-specific overexpression of *Duox* in *mxc*^*mbn1*^. (A-D) The DAPI-stained LGs from mature 3rd instar larvae of control male (*w/Y; He>GFP*) (A), and control larvae harbouring hemocyte-specific overexpression of *Duox* (*w/Y; He>Duox*) (B), *mxc*^*mbn1*^ (*mxc*^*mbn1*^*/Y; He>GFP*) (C), and the mutant larvae harbouring hemocyte-specific overexpression of *Duox* (*mxc*^*mbn1*^*/Y; He>Duox*) (D). Bar; 200 µm. (E) A quantification of the LGs in *mxc*^*mbn1*^ larvae harbouring hemocyte-specific overexpression of *Duox*. For statistical analysis of the differences of the LG size between in *mxc*^*mbn1*^ larvae and in the mutant harbouring hemocyte-specific overexpression, one-way ANOVA with Scheffe’s multiple comparison test was performed (\*\**p*<0.01, n.s.: not significant, n>20). Red lines and error bars represent average value and standard error of mean, respectively.

### Down-regulation of *moladietz*, encoding a maturation factor for Duox in the hemocytes, inhibited the innate signalling pathway and enhanced LG hyperplasia in *mxc*^*mbn1*^ larvae

Numb-interacting protein (NIP), encoded by *moladietz* (*mol*), is a known regulatory factor of Duox in *Drosophila* (Xie et al., 2010). We investigated whether *mol* mRNA level increased in hemocytes of *mxc*^*mbn1*^ larvae using a *mol-lacZ* reporter. Weak anti-β-galactosidase immunostaining signal was detected in the control hemocytes (*w*^*1118*^*/Y; mol-lacZ/+*) (Fig. S4A’). In contrast, a stronger signal was observed in the mutant cells (Fig. S4B’). *mol* expression was significantly higher in *mxc*^*mbn1*^ hemocytes (median 541.43, n = 742) than in control cells (median 472.58, n=1148) (*p* < 0.0001) (Fig. S4C). Thus, *mol* expression was enhanced in the hemocytes of *mxc*^*mbn1*^ larvae. Moreover, the mRNA level of the gene in the mutant larvae was 5 times higher than the level in control larvae at the same developmental stage (Fig. S5D). Thus, *mol* was up-regulated in the LG tumor mutant larvae.

Next, we investigated whether the up-regulation of the gene in hemocytes was required for activation of the innate immune pathway in the mutant fat body. We performed qRT-PCR to quantify the mRNA levels of *Drs* and *Dpt* using total RNAs prepared from fat body of *mxc*^*mbn1*^ larvae harbouring *mol* depletion in the hemocytes (*mxc*^*mbn1*^*/Y; He>molRNAi*) (Fig. S5A). The mRNA level of *Drs*, but not that of *Dpt*, was reduced by 46.3% in *mxc*^*mbn1*^ larvae harbouring hemocyte-specific depletion of *mol*, compared to that in the mutant larvae without the depletion (Fig. S5B). These results suggested that activation of Toll-mediated innate immune pathway in the fat body was suppressed by hemocyte-specific depletion of the Duox activator.

Next, we investigated whether *mol* depletion in the hemocytes affected LG hyperplasia in the *mxc*^*mbn1*^ larvae. We quantified LG size in *mxc*^*mbn1*^ larvae harbouring the hemocyte-specific *mol* depletion (*mxc*^*mbn1*^*/Y; He>molRNAi*) and compared them with those in mutant larvae with control depletion (*mxc*^*mbn1*^*/Y; He>GFPRNAi*). Significant enhancement of LG hyperplasia was observed in *mxc*^*mbn1*^ larvae harbouring the depletion (average LG size was 0.25 mm^2^, n = 24), compared to the average size (0.19 mm^2^, n = 22) in *mxc*^*mbn1*^ larvae with control depletion (*p*<0.01) (Fig. S5C-G). Collectively, up-regulation of the maturation factor of *Duox* also activated the Toll-mediated innate signalling pathway to suppress LG hyperplasia. These results are consistent with the findings that Duox in hemocytes was required for activation of the innate immune pathway in the *mxc*^*mbn1*^ fat body.

### A considerable number of circulating hemocytes were associated with the LG tumors in *mxc*^*mbn1*^ larvae

A previous study has reported that hemocytes containing AMPs secreted from the fat body were attached to the LG tumors in *mxc*^*mbn1*^ larvae (Araki et al., 2019). Therefore, we hypothesized that circulating hemocytes recognized the LG tumors in *mxc*^*mbn1*^ larvae. To confirm this, we investigated whether the hemocytes localized to the mutant LGs. As it was difficult to distinguish the circulating hemocytes and LG cells, we investigated whether normal hemocytes transplanted from normal larvae were recruited to the LG tumors. We collected 0.8 μL hemolymph containing normal hemocytes expressing RFP from control larvae (*w; He>RFP*) and transplanted them into control (*w*^*1118*^*/Y*) and *mxc*^*mbn1*^ larvae at the 3^rd^ instar stage (*mxc*^*mbn1*^*/Y*). Twenty hours after the transplantation, we counted 25.5 RFP^+^ cells (median)/mm^2^ of LG (median) in the *mxc*^*mbn1*^ larvae (Fig. 6B, C), whereas we did not observe any RFP^+^ hemocytes on the LGs in normal larvae (n = 12) (Fig. 6A, C). These genetic data indicated that normal mature hemocytes can be recruited to the LG tumors in *mxc*^*mbn1*^ larvae.

**Fig. 6.**
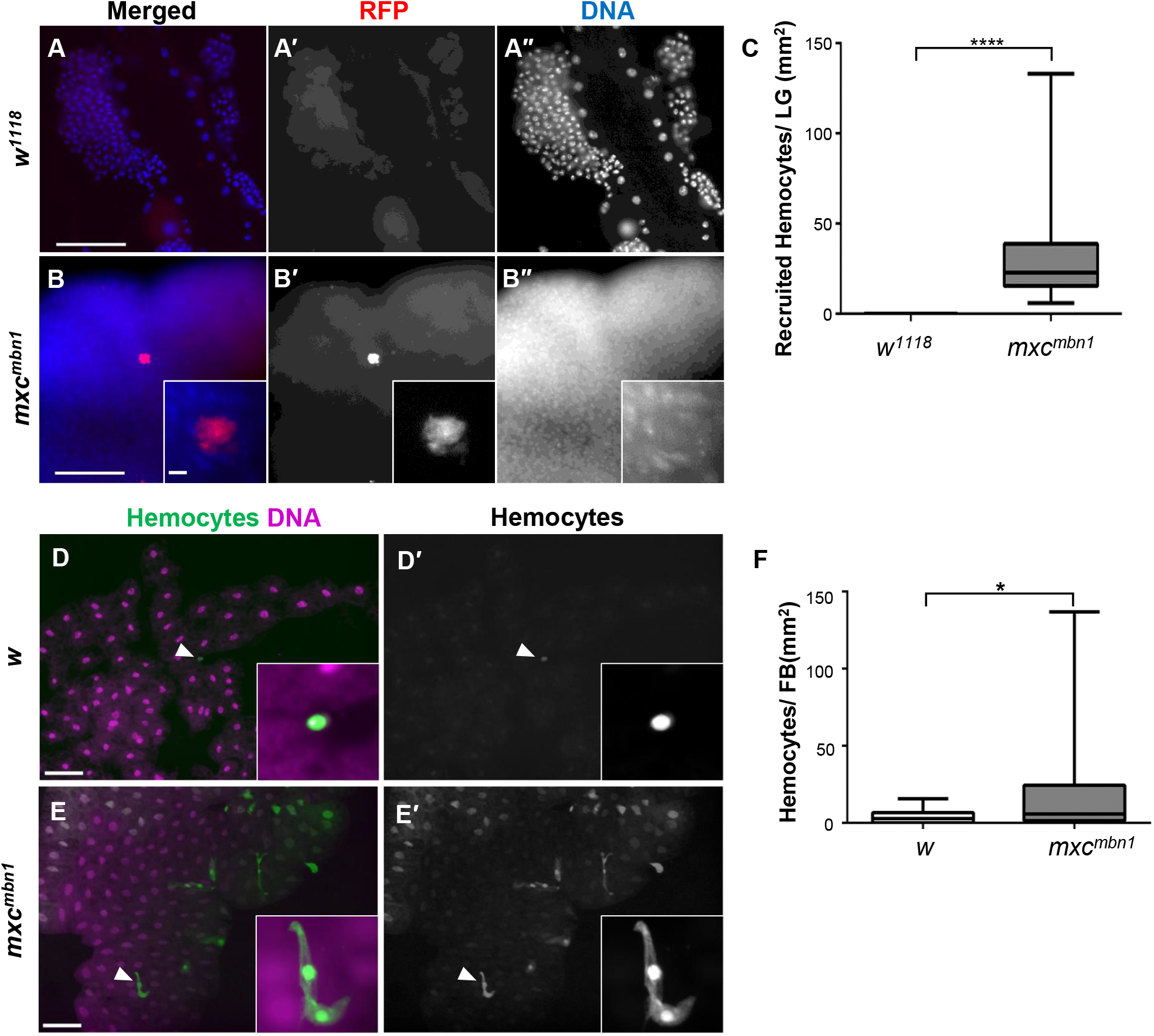
Circulating hemocytes were associated with the LG tumors and fat body in *mxc*^*mbn1*^ larvae. (A, B) Fluorescence image of DAPI-stained LGs from 3^rd^ instar larvae at mature stage, in which RFP-labelled normal hemocytes were transplanted. (A) Normal control LG consisting of the 1^st^ and 2^nd^ lobes (*w*^*1118*^*/Y*). (B) The most anterior region of the LG, corresponding to one-fifth of the whole LG in the *mxc*^*mbn1*^ larva (*mxc*^*mbn1*^*/Y*). (Inset in B) Higher magnification image showing hemocytes on LG. The hemocyte nuclei stained with DAPI (B’) were localized on a focal plane that differed from that in which the fat body nuclei were localized, suggesting that the cells were attached to the surface of the LG. RFP fluorescence, red; DAPI staining; blue. Bar, 100 μm. (C) Quantification of RFP-positive hemocytes observed on the LG. Average numbers of hemocytes on each LG were converted to the number of cells per unit area of the tissue (mm^2^). The Mann-Whitney U test was performed for statistical analysis of the differences in the LG size (\*\*\*\**p* < 0.0001, n ≥ 12). (D, E) Fluorescence images of DAPI-stained fat body and GFPnls-labelled circulating hemocytes in the 3^rd^ instar larvae. (D) Normal control larva (*w/Y; He>GFPnls*). (E) *mxc*^*mbn1*^ larva (*mxc*^*mbn1*^*/Y; He>GFPnls*). (D’, E’) GFP fluorescence in circulating hemocytes harbouring hemocyte-specific expression of GFPnls. (Insets) highly magnified images of a hemocyte (D) and two hemocytes (E) on fat body, shown using arrowheads. Magenta, DAPI staining. Bar, 100 μm. (F) Average numbers of hemocytes on each LG were converted to the number of cells per unit area of the tissue (mm^2^). The Mann-Whitney U test was performed for statistical analysis of the differences in LG size (\**p* < 0.05, n ≥ 26).

### More normal hemocytes were recruited to the fat body in *mxc*^*mbn1*^ larvae than in control larvae

A previous study has reported that several types of AMPs are synthesized and secreted from the fat body in *mxc*^*mbn1*^ larvae (Araki et al., 2019). We hypothesized that circulating hemocytes were involved in transmission of information regarding the existence of tumor cells. To test this, we first investigated whether the hemocytes were localized on the fat body in *mxc*^*mbn1*^ larvae. We scored 2.81 hemocytes (median)/mm^2^ of fat body in the control (*w/Y; He>GFPnls*) (n = 26) (Fig. 6D, F). In contrast, 5.85 GFPnls-labelled cells (median)/mm^2^ were scored in the fat body of the *mxc*^*mbn1*^ larvae (*mxc*^*mbn1*^*/Y; He>GFPnls*) (n = 54), indicating that two times more hemocytes were attached to the LG tumors (*p* < 0.05) (Fig. 6E, F). Thus, the hemocytes in hemolymph were recruited to the LG tumors in *mxc*^*mbn1*^ larvae.

### Up-regulation of *Mmp* genes in *mxc*^*mbn1*^ larvae

Next, we investigated the mechanism via which the hyperplasic LG tumor were recognized and the information was propagated to the fat body. In the other *Drosophila* tumor mutants, ECM in the hypertrophic imaginal discs was decomposed by MMP (Diwanji and Bergmann, 2020, Pastor-Pareja et al., 2008). We hypothesized that MMPs were expressed ectopically in the LG tumors, where they decomposed ECM, and that the hemocytes that recognized the matrix pieces assembled onto the dysplastic sites. To verify this, we examined whether MMP genes were expressed in the mutant LG. *Drosophila* expresses MMP1 and MMP2. Anti-MMP1 immunostaining revealed that the protein was expressed in the cortical zone (CZ) of the LG (Fig. 7A, A”). As the homogenous immunostaining signal was not observed in all cells of the mutant LGs, we measured the MMP1-positive regions of the mutant LGs (Fig. 7B, B”,C). The region was restricted to 8.0% (median, n = 24) of the entire region in control LG. In contrast, the proportion of the regions increased by 26.2% (median, n = 28) and it increased significantly in the *mxc*^*mbn1*^ mutant LGs (Fig. 7C) (*p* < 0.0001).

**Fig. 7.**
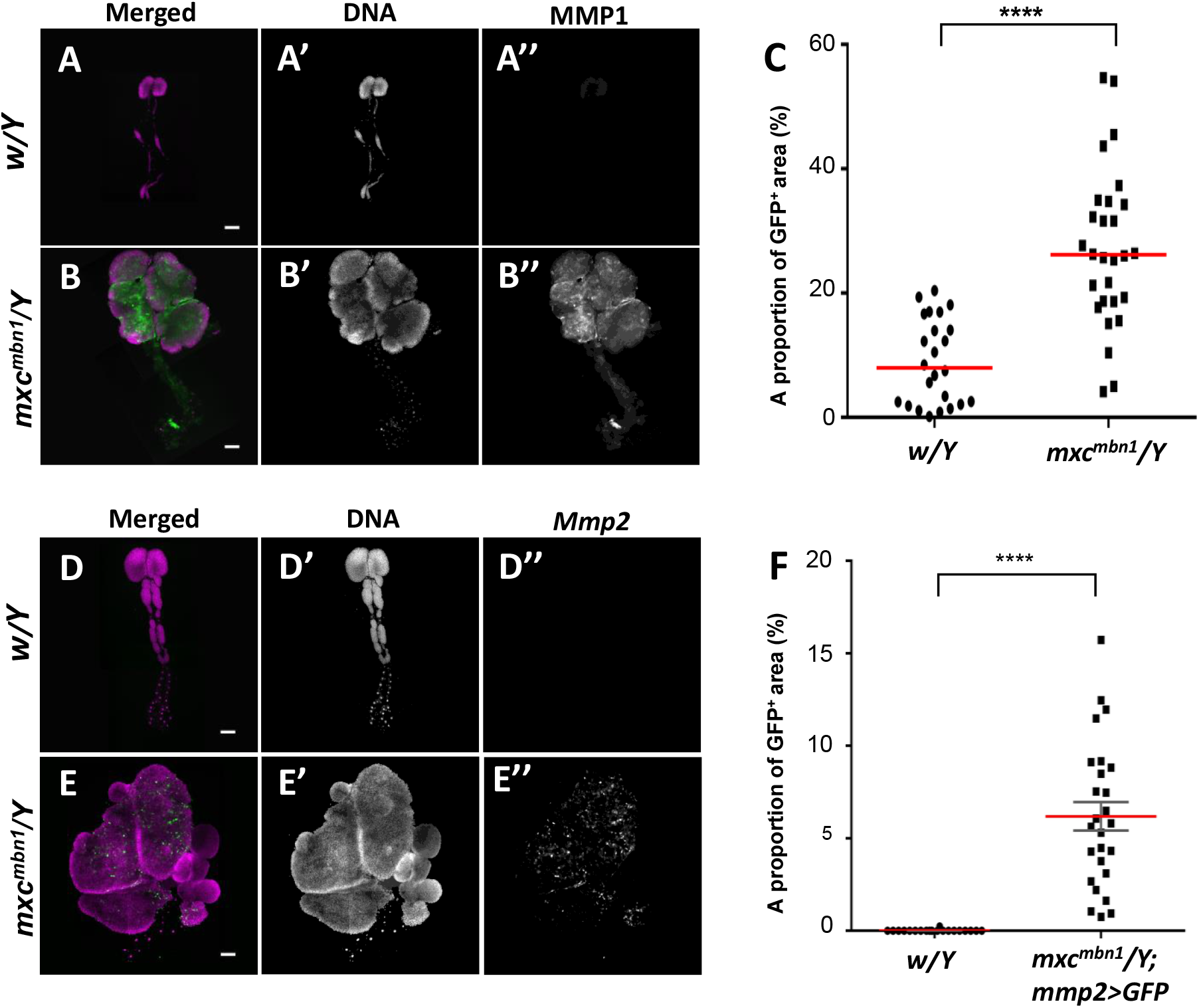
A higher expression of two Matrix metalloproteinases (MMPs) in LGs of *mxc*^*mbn1*^ larvae. (A, B) Anti-MMP1immunostaining of whole LGs from 3^rd^ instar larvae; (A) normal control larva (*w*^*1118*^*/Y*), and (B) *mxc*^*mbn1*^ larva (*mxc*^*mbn1*^*/Y*). Anti-MMPI immunostaining: green (white in A’, B’). DAPI staining: magenta (white in A”, B”). Bar; 100 μm. (C) The proportion of the MMP1-positive region in the LGs from 3^rd^ instar larvae. A percentage of each area per a LG hemisphere was plotted in the graph. For statistical analysis of the differences in the LG size, Mann-Whitney U test was performed (\*\*\*\**p*<0.0001, n>20). (D, E) A visualization of the LG area showing GFP expression under the *Mmp2* enhancer in normal control (*w*^*1118*^*/Y; Mmp2>GFP*) (D), *mxc*^*mbn1*^(*mxc*^*mbn1*^*/Y; Mmp2>GFP*) (E). Bar; 200 μm. GFP fluorescence: green (white in D’, E’). DAPI staining: magenta. (F) The proportion of the region showing *MMP2* gene expression in the LGs from 3^rd^ instar larvae. A percentage of each area per a LG hemisphere was plotted in the graph. For statistical analysis of the differences in the LG size, Mann-Whitney U test was performed (\*\*\*\**p*<0.0001, n≥20). Red lines and error bars represent average value and standard error of mean, respectively.

We next examined whether MMP2 was expressed in the mutant LGs. As antibodies against *Drosophil*a MMP2 are not available, we observed GFP fluorescence in control and *mxc*^*mbn1*^ LGs harbouring *Mmp2-Gal4*, which expressed *Gal4* under the *Mmp2* enhancer. We did not observe any GFP fluorescence in the control LGs (*w/Y;Mmp2>GFP*)(Fig. 7D”), whereas intense fluorescence was observed in the most anterior lobes of the LGs, corresponding to 6.2% of the entire LG region (Fig. 7E”). (Fig. 7F). These observations indicated that MMP2 was expressed ectopically in the mutant LGs.

### Depletion of *Mmp2*, but not that of *Mmp1*, reduced AMP induction in *mxc*^*mbn1*^ larvae

We next investigated how the ectopic expression of *Mmp1* or *Mmp2* influenced AMP expression in the fat body. As dsRNAs against *Mmp1* and *Mmp2* efficiently depleted the relevant mRNAs (Fig. S6A, B), we performed qRT-PCR experiments using total RNAs prepared from the fat body of *mxc*^*mbn1*^ larvae harbouring control depletion (*mxc*^*mbn1*^*/Y; upd3>GFPRNAi*), and *Mpm1* (*mxc*^*mbn1*^*/Y; upd3>Mmp1RNAi*) or *Mpm2* depletion (*mxc*^*mbn1*^*/Y; upd3>Mmp2RNAi*) in the LGs. The levels of *Drs* and *Dpt* mRNAs did not change significantly after the LG-specific *Mmp1* depletion in the *mxc*^*mbn1*^ mutant (Fig. S7A, B middle). In contrast, the mRNA levels of *Drs* and *Dpt* in *mxc*^*mbn1*^ larvae harbouring LG-specific depletion of *Mmp2* decreased by 60.9% and 61.6%, respectively, compared to the levels in the mutant LG with control depletion (Fig. S7A, B left). Thus, down-regulation of *Mmp2*, but not *Mmp1*, influenced activation of the innate immune pathway in the fat body.

### Reduced distribution of basement membrane component in the LGs ectopically expressing *Mmp*2 in *mxc*^*mbn1*^ larvae

We showed that the ectopic expression of *Mmp2* in the mutant LG activated the innate immune pathways. To investigate whether the integrity of basement membrane structure was perturbed in the mutant LGs, we initially visualized the whole LGs from normal 3^rd^ instar larvae harbouring *vkg-GFP* (Fig. 8A, B) under a conventional fluorescence microscope. The collagen IV signal was densely distributed inside the most anterior lobes of the normal LGs (Fig. 8A, 8A’, and inset of 8A’). Intense signals were also observed along the dorsal vessel. In contrast, considerably less signals, some of which ran along the outer periphery of the hyperplasic lobes, were observed in the mutant LG (Fig. 8B, 8B’, and inset of 8B’), although the strong signal in dorsal vessels did not change. Next, we monitored the basement membrane from the surface to the inside of the LGs using a confocal microscope. The GFP signal was abundant in both the surface (Fig. 8D) and on the inside of the confocal plane at approximately 15 μm depth from the surface of the control LGs (Fig. 8E). In contrast, we observed a gradual decline in the fluorescence intensity in the LGs of *mxc*^*mbn1*^ larvae from surface to the depth. The intensity remarkably decreased further and eventually we failed to detect any fluorescence signal at the middle of the LG (Fig. 8G), although the signals were prominent on the surface (Fig. 8F). Furthermore, we quantified the total fluorescence intensity of Vkg-GFP in all lobes of the LG hemispheres (Fig. 8C). Significant reduction in GFP fluorescence was observed in the mutant LGs, compared to normal controls (n > 20, *p* < 0.0001). On the basis of these results, we concluded that the basement membrane was disassembled or lost in the mutant LG ectopically expressing *Mmp2*.

**Fig. 8.**
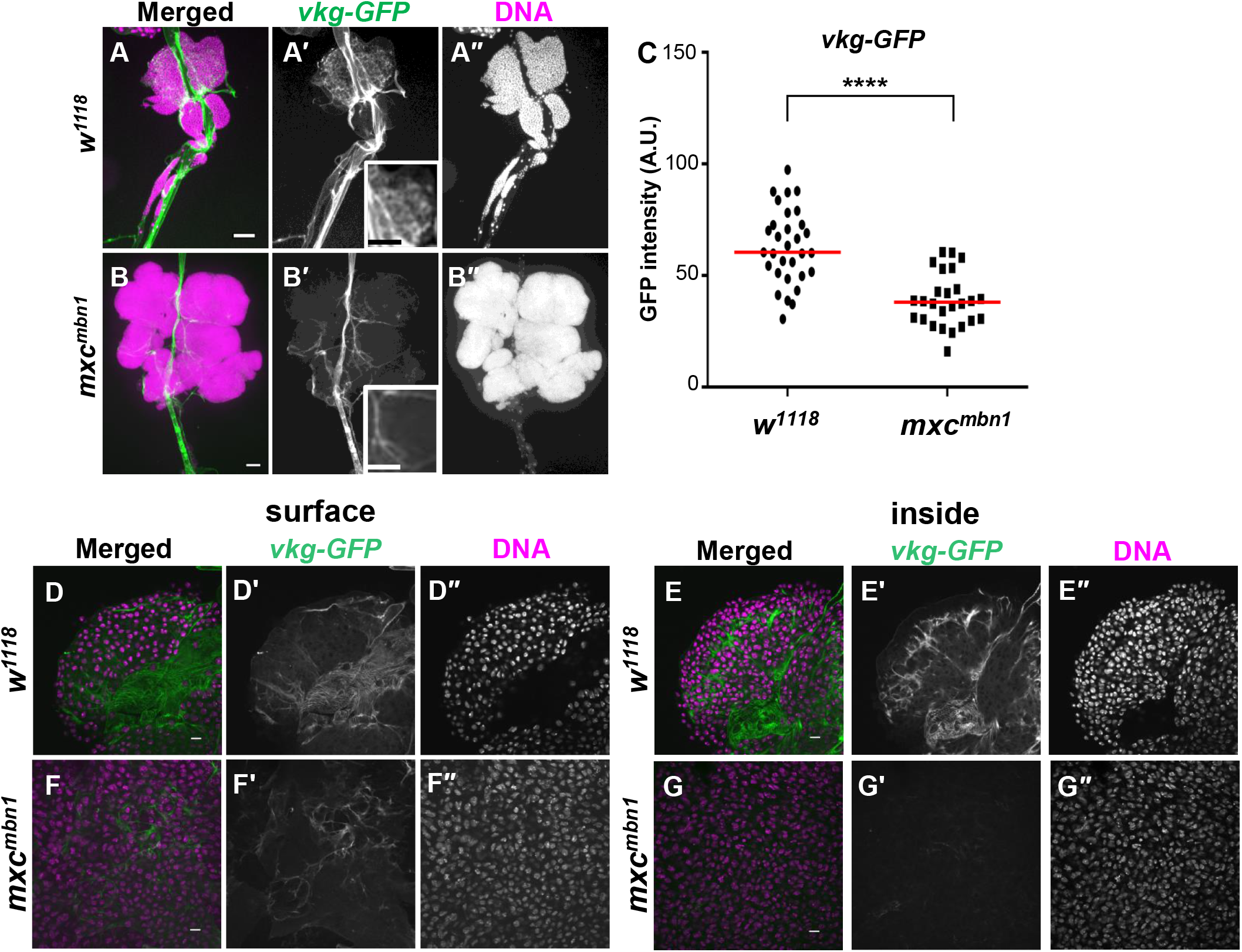
Distribution of the basement membrane component, collagen IV, in whole mount LG from normal and *mxc*^*mbn1*^ larvae. (A, B) Conventional fluorescence micrographs of GFP-tagged collagen IV in each whole LG arranged bilaterally, flanking the dorsal vessel, isolated from 3^rd^ instar control larva (*w/Y; vkg-GFP/+*) (A) and *mxc*^*mbn1*^ larva (B) (*mxc*^*mbn1*^*/Y; vkg-GFP/+*). (C) Arbitrary units of total fluorescence intensity of GFP in lobes in a LG hemisphere (arbitrary unit). For statistical analysis of differences in the LG size, the Mann–Whitney *U* test was performed (\*\*\*\**p*<0.0001, n≥27). Red lines represent median values of the GFP intensity. Scale bars: 100 μm. (D-G) Confocal micrographs of GFP-Collagen IV around LG cells on the surface (C, E) and those inside of the whole mount LGs (F, G) from control (*w/Y; vkg-GFP/+*) (D, F) and *mxc*^*mbn1*^ larvae (*mxc*^*mbn1*^*/Y; vkg-GFP/+*) (E, G). Magenta: DAPI staining. Green: GFP fluorescence.

### Ectopic expression of *Mmp2* in normal wing discs induced *Drs*, a target of the Toll-mediated innate immune signalling pathway

Our genetic data suggested a relationship between the ectopic expression of *Mmp2* in the hyperplastic LGs and activation of the innate immune pathways in the *mxc*^*mbn1*^ larvae. However, determining the mechanism via which the circulating hemocytes recognized the LG tumors was difficult, as the hemocytes originated from tissues. To circumvent this problem, we induced the expression of *Mmp* genes in wing discs and observed the interaction of the hypertrophic tissues with hemocytes. We investigated whether ectopic expression of *Mmp1* or *Mmp2* in the wing discs activated the innate immune signalling pathway in the fat body. Using total RNAs prepared from the fat bodies in control wing discs (*Bx>GFP*), and from those expressing *Mmp1* (*Bx>Mmp1*) and *Mmp2* (*Bx>Mmp2*), we performed qRT-PCR to quantify the mRNA levels of *Drs* (Fig. S8A) and *Dpt* (Fig. S8B). In the larvae harbouring wing disc-specific *Mmp2*, the level of the *Drs* mRNA increased by 190 % of the control level on average, while that of the *Dpt* mRNA decreased by 78 % of that in the control. In contrast, the levels of *Drs* and *Dpt* mRNA in the fat body did not change in larvae harbouring wing disc-specific *Mmp1* expression. These results suggested that ectopic expression of *Mmp2* resulted in activation of the Toll-mediated innate immune pathway, which induced *Drs*.

## DISCUSSION

### Disassembly of basement membrane in the LG tumors ectopically expressing *Mmp2* was involved in tumor recognition in *mxc*^*mbn1*^ larvae

MMPs function as indispensable regulators of cell-cell interactions by controlling the ECM turnover (Page-McCaw et al., 2007). In many human tumors, MMP genes are known to be up-regulated. This evidence suggested that these proteinases are closely related to tumor growth and progression (Egeblad and Werb, 2002). *Drosophila* cells express two types of MMPs, MMP1 and MMP2, during late larval to pupal stages (Page-McCaw et al., 2003). MMP1 preferentially cleaves DE-cadherin to disrupt cell adhesion, for example in the fat body (Jia et al., 2014). In contrast, MMP2 disassembles fat body cells by cleaving basement membrane components (Jia et al., 2014). This is advantageous for the invasion of the tumor cells generated in *Drosophila* imaginal discs (Srivastava et al., 2007; Uhlirova and Bohmann, 2006). In this study, we showed that both MMP1 and MMP2 are highly expressed in the hyperplasic LGs in *mxc*^*mbn1*^ larvae. We demonstrated that MMP2, rather than MMP1 is required for the activation of the innate signalling pathway in response to the LG tumors in the mutant larvae. Indeed, signals of the basement membrane components were reduced or lost in the mutant LGs.

This is because ectopic expression of *Mmp2* enhanced the decomposition of the basement membrane in the mutant larvae, as demonstrated. Furthermore, we observed that the hemocytes were recruited on the LGs in *mxc*^*mbn1*^ larvae (Araki et al., 2019, this study). Another study also reported that the hemocytes were associated with the imaginal disc regions, in which the basement membranes were damaged by JNK-mediated MMP2 from overgrown tissue in *Drosophila* (Diwanji and Bergmann, 2020). Our study further demonstrated that ectopic expression of *Mmp1*, but not *Mmp2* contributes to the activation of the innate immunity pathways in the fat body of *mxc*^*mbn1*^ larvae. This finding also corroborates the observation of a previous report that MMP2, but not MMP1, drives hemocyte recruitment to overgrown imaginal discs (Diwanji and Bergmann, 2020). Our study also demonstrated that the ectopic expression of only *Mmp2* in normal imaginal discs could activate the Toll-mediated pathway, while *Mmp1* does not exhibit any significant effect on the activation of this pathway. Based on these findings, we speculated that the hemocytes the loss of basement membrane integrity in the LG cells possibly contributed to activate the innate immune system in the mutant larvae. Moreover, we observed the disassembly of collagen IV inside the LGs, but not on the surface. This suggests that the ectopic expression of *Mmp2* occurs predominantly in the undifferentiated cells in the medulla zone, rather than in the matured hemocytes in the cortical zone of the LGs.

### Circulating hemocytes were involved in transmission of the information regarding existence of the LG tumors toward the fat body of *mxc*^*mbn1*^ larvae

Our observations suggested that the ectopic expression of *Mmp2* in the LG tumors was required for the activation of the innate immune pathway in the fat body, which was distant from the tumor-bearing tissues. Circulating hemocytes can recognize fragments of ECM generated by MMP ectopically expressed in the tumors (Nastase et al., 2012, Parisi et al., 2014, Diwanji and Bergmann, 2020, this study). In addition to association of the circulating hemocytes with the LG tumors, more hemocytes were also localized on the fat body responsible for the AMP production in the *mxc*^*mbn1*^ larvae. Therefore, we hypothesize that the hemocytes that recognized the tumors moved to the fat body for conveying the information regarding the existence of the tumors toward the innate immunity-responsible tissues. The results of several previous studies support this hypothesis. For example, circulating hemocytes are required to transmit information regarding bacterial infection or local tissue disintegration toward the fat body (Bosch et al., 2019; Parisi et al., 2014; Pastor-Pareja et al., 2008). Another tumor study in *Drosophila* also reported that expression of *spätzle*, encoding a ligand for Toll, was enhanced in hemocytes in *dlg* mutant larvae harbouring tumors. After proteolysis, the active ligand generated could bind and activate the Toll receptor in the fat body (Parisi et al., 2014). In mammals, fragments of ECM disassembled by MMPs induced non-infectious inflammation as damage associated molecular pattern (DAMP), which activated Toll-like receptors as ligands (Nastase et al., 2012). These data support our current hypothesis that basement membrane residues of the tumors produced by MMP2 are recognized by hemocytes, which move toward the fat body and eventually activate innate immune pathway, possibly as the DAMPS in *mxc*^*mbn1*^ larvae.

### ROS generated by active Duox in circulating hemocytes contributed to the activation of the innate immune system in the fat body to induce AMPs

We observed that ROS accumulated in the circulating hemocytes in *mxc*^*mbn1*^ larvae and further demonstrated that elimination of ROS by antioxidants suppressed the activation of Toll- and Imd-mediated innate immune signalling pathways in the fat body of *mxc*^*mbn1*^ larvae. Although ROS are known to activate innate immune pathways, it is unlikely that ROS themselves act as signal messengers via the hemolymph, as ROS are unstable *in vivo* (Dickinson and Chang, 2011). Instead, we found that the circulating hemocytes were associated with the LG tumors, as well as with the fat body in *mxc*^*mbn1*^ larvae. Therefore, we speculated that the hemocytes that recognized the tumor cells might transfer the information toward the fat body. The Toll-mediated pathway was activated via ROS when *Drosophila* larvae were infected by a parasitoid wasp (Louradour et al., 2017). Moreover, the hemocytes that recognized the stimuli activated the Toll-mediated signalling pathway and consequently induced the expression of *Cecropin-C (CecC)* (Chakrabarti and Visweswariah, 2020). In mammalian cells, ROS is consistently required for activation of the IKK complex in the innate immune pathway and for stimulating nuclear import of Relish (Morgan and Liu, 2011). These observations indicated that ROS play an important role in activating innate immune signalling. Tissue disintegration can be recognized by hemocytes, and the information is transmitted to the fat body producing AMPs (Capilla et al., 2017). In this study, we showed that *Duox* was up-regulated in *mxc*^*mbn1*^ larvae. This was indispensable for the activation of the innate immune pathway. Therefore, ROS production and accumulation in circulating hemocytes is required for the activation of the innate immune pathways. Moreover, we demonstrated that a maturation factor for Duox was similarly up-regulated in the mutant hemocytes. This supported the idea that the oxidase was critical for the activation of the innate immune pathway. Consistently, its mammalian orthologue, DUOXA, is also essential for translocating the oxidase to the cell surface. DUOXA ensures efficient ROS production by DUOX, as it associates continuously with the oxidase (Grasberger and Refetoff, 2006; Qin et al., 2004).

Duox and its activator, were up-regulated in the hemocytes of *mxc*^*mbn1*^ larvae, but not in the fat body producing AMPs. This suggested that the hemocytes conveyed the ROS toward the fat body via the hemolymph. ROS is involved in processing of the ligand, Spätzle, which binds to the Toll receptor (Louradour et al., 2017). In larvae harbouring epithelial tumors, Spätzle is also induced and activated via ROS-dependent proteolysis in hemocytes, which subsequently activates the Toll receptor (Parisi et al., 2014). Therefore, we speculate that the expression of *Spätzle* might be induced in the mutant hemocytes and subsequently it might be activated in a ROS-dependent manner in the fat body, as observed in this study. Further investigations are warranted to conclusively establish the mechanism by which the expression of the gene is elevated, and the active ligand is generated.

Considering our results and those of previous studies, we proposed the following model to explain the mechanism via which *Drosophila* LG tumors are detected and eliminated by the innate immune system. Ectopically expressed MMP2 in the LG tumors in *mxc*^*mbn1*^ larvae degrades the basement membrane in the tissue. The hemocytes that recognize the tissue disintegration accumulated on the LG tumors. Consequently, simultaneous induction of *Duox* and *mol*, they produced the oxidase and secreted ROS around the cells. Thereby, the hemocytes producing ROS continuously conveyed the information regarding the existence of tumors toward the fat body. Thereafter, the cells migrated to the fat body and activated the Toll-mediated signalling pathway on the tissue using active Toll ligand generated via ROS-dependent proteolysis. Finally, the AMPs were produced and secreted from the fat body to supress the LG tumors.

In this study, we observed that the disassembly of basement membrane due to the ectopically expressed MMP2 was involved in the tumor recognition by macrophage-like hemocytes in *mxc*^*mbn1*^ larvae in *Drosophila*. Although several studies have demonstrated the ectopic expression of MMPs in epithelial tumors, only a few of them have presented substantial evidence that the disassembly of basement membrane by MMP2 is required for tumor recognition (Beaucher et al., 2007, Kudo et al., 2012). Our previous study established the association between the hemocytes that incorporated AMPs with tumors; however, the mechanism by which the hemocytes recognized the ECM fragments remained elusive. This study revealed that the ROS producing-hemocytes moved to the fat body to activate the innate immune pathways. These findings highlight the mechanism by which macrophage-like *Drosophila* hemocytes also relay the information toward the fat body apart from using ROS and activate the innate immune system. In our future studies, we would like to elucidate the mechanism of activation of the Toll ligand Spätzel on the fat body of *mxc*^*mbn1*^ larvae by the ROS generated from the hemocytes.

## Materials and Methods

### *Drosophila* stocks and their husbandry

Canton S was used as the wild-type stock, and *w*^*1*^ as the normal control stock. A stock carrying the *mxc* lethal mutation showing the LG tumor phenotype (*mxc*^*mbn1*^) were used (Araki et al., 2019; Kurihara et al., 2020; Remillieux-Leschelle et al., 2002). As heterozygotes for the mutation were maintained under a balancer chromosome carrying *sqh-RFP*, the hemizygotes were selected as larvae without RFP expression. For the UAS-dependent expression of mRNAs or dsRNA in hemocytes, we used *P*{*w*^*+mC*^*=He-Gal4*.*Z*}(*He-Gal4*) obtained from Bloomington *Drosophila* Stock Center (BDSC, Indiana University, Bloomington, IN, USA) (Stofanko et al., 2008). To induce the gene expression in immature hemocyte precursors in lymph gland, *P{upd3-GAL4}*(*upd3-Gal4*) (Jung et al., 2005) from N. Perrimon (Harvard Medical School, Boston, MA, USA) was used. *P{w*^*+mW*.*hs*^*= GawB}Bx*^*MS1096*^ (*Bx-Gal4*) (BL8860 from BDSC) was used to induce ectopic gene expression in wing imaginal discs (Capdevila and Guerrero, 1994).

For the ectopic expression of Duox (dDuox), MMP1 and MMP2, we used *UAS-dDuox* (*UAS-Duox*)(Ha et al., 2005), which was a gift from W. Lee (Seoul National Univ., Republic of Korea), *P{w*^*+mC*^*=UAS-Mmp1*.*f1}3* (*UAS-Mmp1*) (BL58701 from BDSC) and *P{w*^*+mC*^*=UAS-Mmp2*.*P}* (*UAS-Mmp2*) (BL58706 from BDSC), respectively. For dsRNA-dependent gene silencing of gene encoding Dual oxidase(Duox), *mol* gene for a Numb-interacting protein(NIP) required for Duox maturation, genes for Mmp1, and Mmp2, *P{TRiP*.*GL00678}attP40* (*UAS-DuoxRNAi*^*GL00678*^) from BDSC (BL38907 from BDSC), *P{TRiP*.*HMS02560}attP40* (*UAS-molRNAi*^*HMS02560*^) (BL42867 from BDSC), *P{KK108894}VIE-260B* (*UAS-Mmp1RNAi*^*KK108894*^) from Vienna Drosophila Resource Center (VDRC) (Vienna, Austria) (#101505), and *P{TRiP*.*HMJ23143}attP40* (*UAS-Mmp2RNAi*) from National Institute of Genetics (Mishima, Japan) (#11605). To monitor gene expression of the *mol* gene, and *Mmp2* gene, *P{lacW}mol*^*k11524a*^ (*mol-lacZ*) (BL12173 from BDSC) and *P{w*^*+mW*.*hs*^*=GawB}Mmp2*^*NP0509*^ (*Mmp2-GAL4*) (Srivastava et al., 2007) (#103625 from DGRC) were used, respectively. To visualize basement membrane component, collagen IV, *PBac{fTRG00595*.*sfGFP-TVPTBF}VK00033* (*Vkg-GF*P)(#318167 from VDRC) was used, respectively. A redox GFP reporter, *P{gstD1-GFP*.*S}* (*gstD1-GFP*) was used to estimate the extent of oxidative stress accumulation in larval tissues (Sykiotis and Bohmann, 2008; Le et al., 2019). This stock was a gift of D. Bohmann (Rochester Univ., Rochester, NY, USA).

All *Drosophila* stocks were maintained on standard cornmeal food, as previously described (Oka et al., 2015). Gal4-dependent expression was measured at 28°C. Other experiments and stock maintenance were conducted at 25°C.

### LG preparation

Normal controls (*+*/Y or *w*/Y) pupated at 6 days (28°C) and 7 days (25°C) after egg laying (AEL), whereas some of the *mxc*^*mbn1*^ mutant remained in 3rd instar larval stage at 10 days (28°C) and 11 days (25°C) AEL (Kurihara et al., 2020). To minimize the possibility of a delay that might allow the tissue to grow, the comparative analysis of hemizygous mutants and controls was performed on the same day (5 days AEL at 28°C), when the wandering 3rd instar larval stage was seen. Alternatively, the tissues were collected from hemizygous mutant larvae one day after the timing of the LG collection from control larvae. For the staging of the larvae, parent flies were transferred into a new culture vial and left there to lay eggs for 24 h. A pair of anterior lobes of the LG without connected cardiac cells from mature stage larvae were isolated and fixed with 3.7% formaldehyde for 5 min. The fixed samples were mildly flattened under constant pressure using an apparatus so that the tissue became spread out into cell layers with a constant thickness. A pair of lobes of the LG without connected cardiac cells from mature stage larvae were isolated and fixed with 3.7% formaldehyde. The fixed each LG pair was mildly flattened under constant pressure so that the tissue became spread out into cell layers with a constant thickness as described (Araki et al., 2019). The microscope images acquired as multiple images were assembled to single images using photoshop (CS6 version, Adobe systems, San Jose, CA, USA). The LG area of each DAPI-stained sample was measured using ImageJ ver.1.47 (https://imagej.nih.gov/ij/).

### LG immunostaining

For immunostaining, LGs were dissected from matured 3rd instar larvae and fixed in 3.7% paraformaldehyde for 15 min. After repeated washing, the fixed samples were incubated with primary antibody at 4°C for overnight. The following anti-Mmp1 antibodies (#3A6B4, #3B8D12, and #5H7B11) were mixed and used (1:100 for each; DSHB, IA, USA). After extensive washing, specimens were incubated with Alexa 594 secondary antibody (1:400; Molecular Probe, USA). The LG specimens were observed under a fluorescence microscope (Olympus, Tokyo, Japan, model: IX81), outfitted with excitation, emission filter wheels (Olympus). The fluorescence signals were collected using a 10x dry objective lens. Specimens were illuminated with UV filtered and shuttered light using the appropriate filter wheel combinations through a GFP filter cube. GFP fluorescence images were captured with a CCD camera (Hamamatsu Photonics, Shizuoka, Japan). Image acquisition was controlled through the Metamorph software version 7.6 (Molecular Devices, Sunnyvale, CA, USA) and processed with Adobe Photoshop CS. The basement membrane of the LG cells was observed under a confocal microscope from the surface to the inside of the tissue (Fv10i, Olympus, Tokyo, Japan) by altering the focus along the z-axis. The confocal images obtained were then processed by the Fv10i software and Adobe photoshop CS (Adobe KK, Tokyo, Japan).

### Immunostaining of hemocytes in larval hemolymph

Single mature larvae at the 3rd instar stage were dissected in Drosophila Ringer solution (DR) (10 mM, pH 7.2, Tris-HCl, 3 mM CaCl2·2H2O, 182 mM KCL, 46 mM NaCl) on a slide glass so as to allow circulating hemocytes to release into the DR outside the larvae. Whole aliquots of the cell suspension were collected as much as possible, and hemocytes in the cell suspension were counted using a hemocytometer (Araki et al., 2019). The cells were fixed in 4% paraformaldehyde for 10 min after placing the small amount of DR containing circulating hemocytes on a slide glass and leaving for evaporating. We used anti-β-galactosidase antibody (MP Biomedicals (Irvine, CA, USA), #02150039, 1:2000), anti-P1 antibody for plasmatocytes (Kurucz et al., 2007, 1:100) which was a gift from I. Ando (Hungarian Academy of Sciences, Budapest, Hungary) as primary antibodies. The number of circulating hemocytes in the fluorescence microscope images was counted by Image J.

For monitoring reactive oxygen species (ROS) in hemocytes and larval tissues, dihydroethidium (DHE) (#181094, Life Technologies (Carlsbad, CA, USA)) was used. We quantified the intensity levels of the DHE fluorescence in hemocytes, the GFP fluorescence in the hemocytes of *gstD-GFP* larvae, and anti-β-galactosidase immunostaining signal in *mol-lacZ* using Image J. The cells were classified into five categories according to the intensity levels: cells showing less than 4,000 intensity values in ImageJ were of background level or below category (I), those with 4,000 to 20,000 intensity values were in category II, with 20,000 to 40,000 intensity values in moderate category (III), with 40,000 to 60,000 intensity values in intense category (IV), and higher than 60,000 intensity values in the more intense category (V). We classified the GFP-positive cells in *gstD-GFP* larvae into four groups according to the fluorescence intensity; cells showing less than 30 values were in category I, 30 to 70 in category II, 70 to 150 in III, and more than 150 in IV.

### 5. qRT-PCR analysis

Total RNA was extracted from whole larvae at 3rd instar stage, and larval fat body, guts, LGs and hemocytes in hemolymph with each genotype using the Trizol reagent (Invitrogen, Carlsbad, CA, USA). cDNA synthesis from the total RNA was carried out using the PrimeScriptTM High Fidelity RT-PCR Kit (TaKaRa, Shiga, Japan) using an oligo dT primer. Real-time PCR was performed using the FastStart Essential DNA Green Master (Roche, Mannheim, Germany) and a Light Cycler Nano instrument (Roche, Mannheim, Germany). According to a software the Primer3Plus (http://www.bioinformatics.nl/cgi-bin/primer3plus.cgi), the following primers were synthesized:

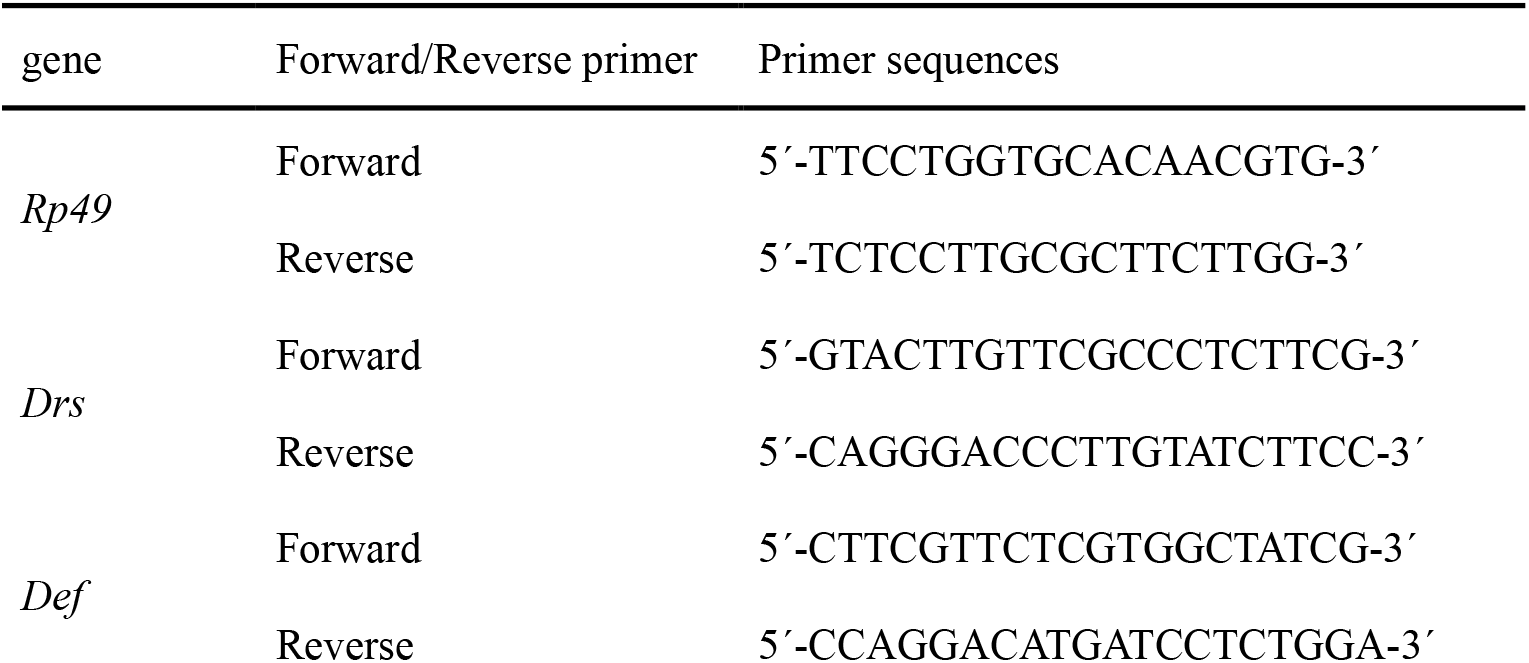
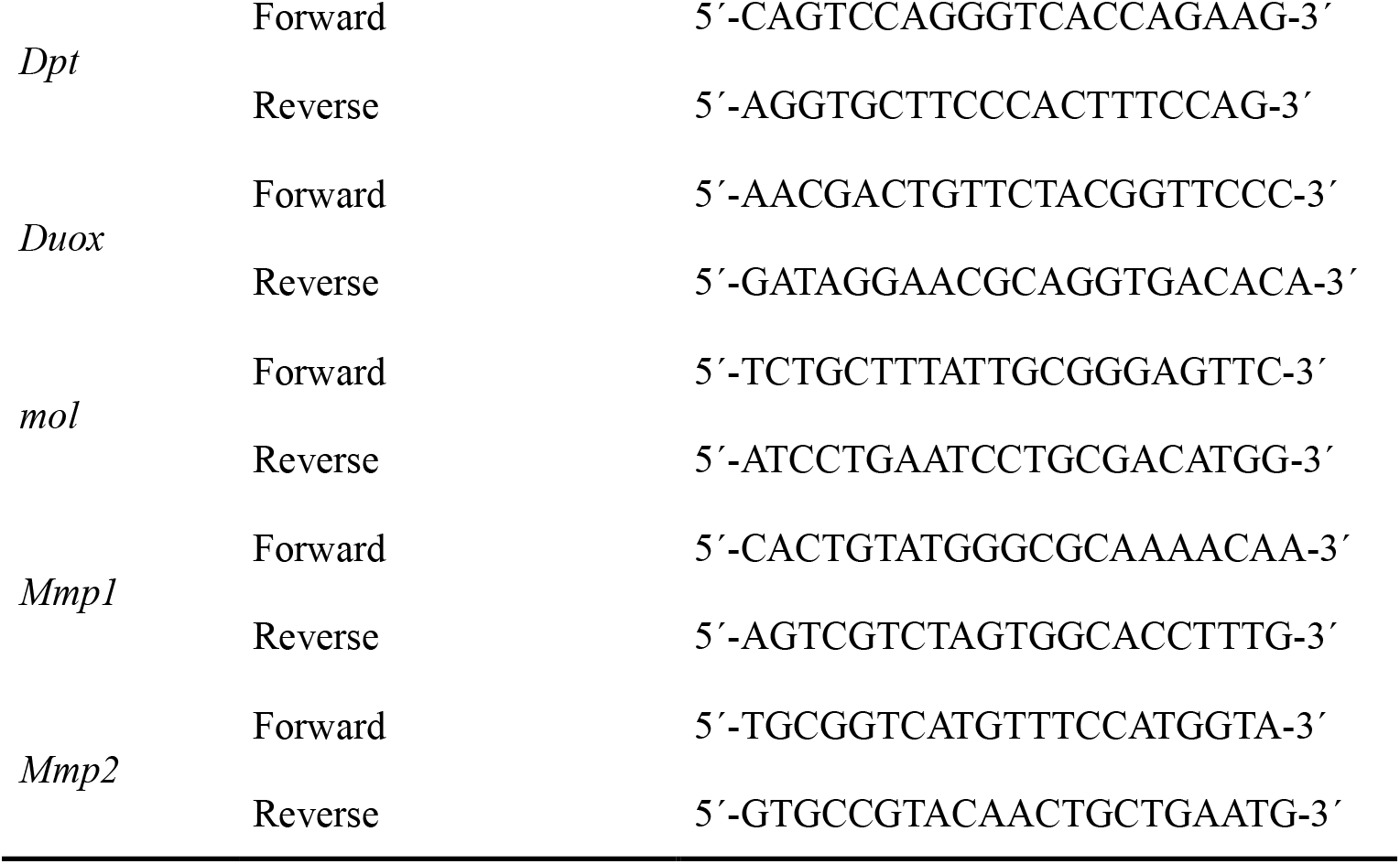

### Transplantation of hemocytes in *Drosophila* larvae

A small aliquot (0.8 μl) of larval hemolymph containing hemocytes (2,151±301 cells on average) or pieces of the LG were performed using a glass needle into a recipient 3^rd^ instar larva. The needles were generated from G1.2 (Narishige Co., Tokyo, Japan) using a puller, PN-31, and used after sharpening of the tip. The donor hemocytes were injected within 5min after dissection of recipient larvae to avoid melanization and clogging. The injected larvae were put on wet blocking papers for 1 hr to recover them from the damage and raised on a standard food for 20 h before observation.

### Feeding experiments of the antioxidant

30 embryos laid within a day were transferred to a standard food supplemented with 0.1 mg/ml N-acetyl cysteine(NAC) (Sigma-Aldrich(Saint Louis, MO, USA),#A7250) and allowed the larvae to feed the food at 25°C.

### Statistical analysis

Results of the LG area measurements were presented as scatter plots created using GraphPad Prism 6 or Microsoft Office Excel 2016 (Microsoft (Redmond, WA, USA)). The area in pixels was calculated and an average value was determined for each LG. Each single dataset was assessed using Welch’s t test or Student’s t test as described previously (Araki et al., 2019; Kurihara et al., 2020). The F -test was performed to determine equal or unequal variance. *p*-values were calculated using Welch’s t test of unequal variance if the value was less than 0.05. *p*-values were calculated using the Student’s t test of equal variance when the F -value was greater than 0.05. The statistical significance is described in each figure legend: *;*p* < 0.05, **; *p* < 0.01, ***; *p* < 0.001, and ****; *p* < 0.0001.

## Acknowledgements

We appreciate Mayo Araki (Kyoto Institute of Technology, Kyoto, Japan) for her dedicated efforts at the initial steps of this study. We also acknowledge for N. Perrimon, W. Lee and D. Bohman for providing fly stocks and I. Ando for P1 antibody. We also thank Bloomington Stock Centre, Kyoto Stock Centre, National Institute of Genetics and Vienna *Drosophila* Resource Centre for providing fly stocks.

## Author contributions

SK carried out most of observations of the LG phenotypes, immunostaining experiments of larval tissues, and qRT-PCR experiments. KT performed confocal observation of the LGs. YHI planned, organized the project, and led the interpretation of the data. YHI wrote the manuscript. All authors read and approved the final manuscript.

## REFERENCES

Araki, M., Kurihara, M., Kinoshita, S., Awane, R., Sato, T., Ohkawa, Y., and Inoue, Y.H. (2019). Anti-tumor effects of antimicrobial peptides, components of the innate immune system, against haematopoietic tumors in Drosophila mxc mutants. Dis. Model. Mech. 12, dmm037721.

Arefin B, Kunc M, Krautz R, Theopold U. (2017). The Immune phenotype of three Drosophila leukemia models. G3 (Bethesda). 7, 2139–2149.

Beaucher, M., Hersperger, E., Page-McCaw, A., and Shearn, A. (2007). Metastatic ability of Drosophila tumors depends on MMP activity. Dev Biol. 303, 625–634.

Bilder, D., Ong, K., His, T.C., Adiga, K., and Kim, J. (2021). Tumour-host interactions through the lens of Drosophila. Nat Rev Cancer. 21, 687–700.

Bosch, P.S., Makhijani, K., Herboso, L., Gold, K.S., Baginsky, R., Woodcock, K.J., Alexander, B., Kukar, K., Corcoran, S., Jacobs, T., Ouyang, D., Wong, C., Ramond, E.J.V., Rhiner, C., Moreno, E., Lemaitre, B., Geissmann, F., and Brückner, K. (2019). Adult Drosophila lack hematopoiesis but rely on a blood cell reservoir at the respiratory epithelia to relay infection signals to surrounding tissues. Dev. Cell 51, 787–803.

Brennan, C.A., and Anderson, K.V. (2004). Drosophila: The genetics of innate immune recognition and response. Annu. Rev. Immunol. 22, 457–483.

Buchon, N., Silverman, N. and Cherry, S. (2014). Immunity in Drosophila melanogaster--from microbial recognition to whole-organism physiology. Nat. Rev. Immunol. 14, 796–810.

Capdevila, J., and Guerrero, I. (1994). Targeted expression of the signaling molecule decapentaplegic induces pattern duplications and growth alterations in Drosophila wings. EMBO J. 13, 4459–4468.

Capilla, A., Karachentsev, D., Patterson, R.A., Hermann, A., Juarez, M.T., and McGinnis, W. (2017). Toll pathway is required for wound-induced expression of barrier repair genes in the Drosophila epidermis. Proc. Natl. Acad. Sci. U S A 114, 2682–2688.

Chakrabarti, S., and Visweswariah, S.S. (2020). Intramacrophage ROS primes the innate immune system via JAK/STAT and Toll activation. Cell Rep. 33, 108368.

Choe, K.-M., Lee, H., and Anderson, K.V. (2005). Drosophila peptidoglycan recognition protein LC (PGRP-LC) acts as a signal-transducing innate immune receptor. Proc. Natl. Acad. Sci. U S A 102, 1122–1126.

Dickinson, B.C., and Chang, C.J. (2011). Chemistry and biology of reactive oxygen species in signaling or stress responses. Nat. Chem. Biol. 7, 504–511.

Diwanji, N., and Bergmann, A. (2020). Basement membrane damage by ROS- and JNK-mediated Mmp2 activation drives macrophage recruitment to overgrown tissue. Nat. Commun. 11, 3631.

Egeblad, M., and Werb, Z. (2002). New functions for the matrix metalloproteinases in cancer progression. Nat. Rev. Cancer 2, 161–174.

Evans, C.J., Hartenstein, V., and Banerjee, U. (2003). Thicker than blood: conserved mechanisms in Drosophila and vertebrate hematopoiesis. Dev. Cell 5, 673–690.

Fehlbaum P, Bulet P, Michaut L, Lagueux M, Broekaert WF, Hetru C, Hoffmann JA. (1994). Insect immunity. Septic injury of Drosophila induces the synthesis of a potent antifungal peptide with sequence homology to plant antifungal peptides. J Biol Chem. 269, 33159–33163.

Georgel, P., Naitza, S., Kappler, C., Ferrandon, D., Zachary, D., Swimmer, C., Kopczynski, C., Duyk, G., Reichhart, J.M., and Hoffmann, J.A. (2001). Drosophila immune deficiency (IMD) is a death domain protein that activates antibacterial defense and can promote apoptosis. Dev. Cell 1, 503–514.

Gobin, E., Bagwell, K., Wagner, J., Mysona, D., Sandirasegarane, S., Smith, N., Bai, S., Sharma, A., Schleifer, R., She, J.X. (2019). A pan-cancer perspective of matrix metalloproteases (MMP) gene expression profile and their diagnostic/prognostic potential. BMC Cancer. 19, 581.

Gottar, M., Gobert, V., Michel, T., Belvin, M., Duyk, G., Hoffmann, J.A., Ferrandon, D., Royet, J. (2002). The Drosophila immune response against Gram-negative bacteria is mediated by a peptidoglycan recognition protein. Nature. 416, 640–644.

Govind, S. (2008). Innate immunity in Drosophila: Pathogens and pathways. Insect Sci. 15, 29–43.

Grasberger, H., and Refetoff, S. (2006). Identification of the maturation factor for dual oxidase. Evolution of an eukaryotic operon equivalent. J. Biol. Chem. 281, 18269–18272.

Ha, E.-M., Oh, C.-T., Bae, Y.S., and Lee, W.-J. (2005). A direct role for dual oxidase in Drosophila gut immunity. Science 310, 847–850.

Hoffmann JA. (2003). The immune response of Drosophila. Nature. 426, 33–38.

Hoffmann, J.A., and Reichhart, J.M. (2002). Drosophila innate immunity: an evolutionary perspective. Nat. Immunol. 3, 121–126.

Jang, I.H., Chosa, N., Kim, S.H., Nam, H.J., Lemaitre, B., Ochiai, M., Kambris, Z., Brun, S., Hashimoto, C., Ashida, M., Brey, P.T., and Lee, W.J. (2006). A Spätzle-processing enzyme required for toll signaling activation in Drosophila innate immunity. Dev Cell. 10, 45–55.

Jia, Q., Liu, Y., Liu, H., and Li, S. (2014). Mmp1 and Mmp2 cooperatively induce Drosophila fat body cell dissociation with distinct roles. Sci. Rep. 4, 7535.

Jung, S.H., Evans, C.J., Uemura, C., and Banerjee, U. (2005). The Drosophila lymph gland as a developmental model of hematopoiesis. Development 132, 2521–2533.

Kalamarz, M.E., Paddibhatla, I., Nadar, C., Govind, S. (2012). Sumoylation is tumor-suppressive and confers proliferative quiescence to hematopoietic progenitors in Drosophila melanogaster larvae. Biol Open. 1, 161–172.

Kessenbrock, K., Plaks, V., and Werb, Z. (2010). Matrix metalloproteinases: regulators of the tumor microenvironment. Cell. 141, 52–67.

Kim, M.J., and Choe, K.M. (2014). Basement membrane and cell integrity of self-tissues in maintaining Drosophila immunological tolerance. PLoS Genet. 10, e1004683.

Kleino, A., Valanne, S., Ulvila, J., Kallio, J., Myllymäki, H., Enwald, H., Stöven, S., Poidevin, M., Ueda, R., Hultmark, D., Lemaitre, B., and Rämet, M. (2005). Inhibitor of apoptosis 2 and TAK1-binding protein are components of the Drosophila Imd pathway. Embo J. 24, 3423–3434.

Kudo Y, Iizuka S, Yoshida M, Tsunematsu T, Kondo T, Subarnbhesaj A, Deraz EM, Siriwardena SB, Tahara H, Ishimaru N, Ogawa I, and Takata T. (2012). Matrix metalloproteinase-13 (MMP-13) directly and indirectly promotes tumor angiogenesis. J Biol Chem. 287, 38716–3828.

Kurihara, M., Komatsu, K., Awane, R., and Inoue, Y.H. (2020). Loss of Histone Locus Bodies in the Mature Haemocytes of Larval Lymph Gland Result in Hyperplasia of the Tissue in mxc Mutants of Drosophila. Int. J. Mol. Sci. 21, 1586.

Kurihara, M., Takarada, K., and Inoue, Y.H. (2020). Enhancement of leukemia-like phenotypes in Drosophila mxc mutant larvae due to activation of the RAS-MAP kinase cascade possibly via down-regulation of DE-cadherin. Genes to Cells 25, 757–769.

Kurucz, E., Márkus, R., Zsámboki, J., Folkl-Medzihradszky, K., Darula, Z., Vilmos, P., Udvardy, A., Krausz, I., Lukacsovich, T., Gateff, E., Zettervall, C.J., Hultmark, D., and Andó, I. (2007). Nimrod, a putative phagocytosis receptor with EGF repeats in Drosophila plasmatocytes. Curr. Biol. 17, 649–654.

Lanot, R., Zachary, D., Holder, F., and Meister, M. (2001). Postembryonic hematopoiesis in Drosophila. Dev. Biol. 230, 243–257.

Le, T.D., Nakahara, Y., Ueda, M., Okumura, K., Hirai, J., Sato, Y., Takemoto, D., Tomimori, N., Ono, Y., Nakai, M., Shibata, H., and Inoue, Y.H. (2019). Sesamin suppresses aging phenotypes in adult muscular and nervous systems and intestines in a Drosophila senescence-accelerated model. Eur Rev Med Pharmacol Sci. 23, 1826–1839.

Lemaitre B, Hoffmann J. A. (2007) The host defense of Drosophila melanogaster. Annu. Rev. Immunol. 25, 697–743.

Lemaitre, B., Kromer-Metzger, E., Michaut, L., Nicolas, E., Meister, M., Georgel, P., Reichhart, J.M., and Hoffmann, J.A. (1995). A recessive mutation, immune deficiency (imd), defines two distinct control pathways in the Drosophila host defense. Proc. Natl. Acad. Sci. U S A 92, 9465–9469.

Lemaitre B, Reichhart JM, Hoffmann JA. (1997). Drosophila host defense: differential induction of antimicrobial peptide genes after infection by various classes of microorganisms. Proc Natl Acad Sci U S A. 94, 14614–14619.

Louradour, I., Sharma, A., Morin-Poulard, I., Letourneau, M., Vincent, A., Crozatier, M., and Vanzo, N. (2017). Reactive oxygen species-dependent Toll/NF-κB activation in the Drosophila hematopoietic niche confers resistance to wasp parasitism. Elife 6, e25496.

Martinelli, C., and Reichhart, J.M. (2005). Evolution and integration of innate immune systems from fruit flies to man: lessons and questions. J. Endotoxin Res. 11, 243–248.

Morgan, M.J., and Liu, Z.-g. (2011). Crosstalk of reactive oxygen species and NF-κB signaling. Cell Res. 21, 103–115.

Morisato, D., and Anderson, K.V. (1994). The spätzle gene encodes a component of the extracellular signaling pathway establishing the dorsal-ventral pattern of the Drosophila embryo. Cell 76, 677–688.

Nastase, M.V., Young, M.F., and Schaefer, L. (2012). Biglycan: a multivalent proteoglycan providing structure and signals. J. Histochem. Cytochem. 60, 963–975.

Oka, S., Hirai, J., Yasukawa, T., Nakahara, Y., and Inoue, Y.H. (2015). A correlation of reactive oxygen species accumulation by depletion of superoxide dismutases with age-dependent impairment in the nervous system and muscles of Drosophila adults. Biogerontology 16, 485–501.

Page-McCaw, A., Ewald, A.J., Werb, Z. (2007). Matrix metalloproteinases and the regulation of tissue remodelling. Nat. Rev. Mol. Cell Biol. 2007 8, 221–233.

Page-McCaw, A., Serano, J., Santé, J.M., and Rubin, G.M. (2003). Drosophila matrix metalloproteinases are required for tissue remodeling, but not embryonic development. Dev. Cell 4, 95–106.

Parisi, F., Stefanatos, R.K., Strathdee, K., Yu, Y., and Vidal, M. (2014). Transformed epithelia trigger non-tissue-autonomous tumor suppressor response by adipocytes via activation of Toll and Eiger/TNF signaling. Cell Rep. 6, 855–867.

Parvy, J.P., Yu, Y., Dostalova, A., Kondo, S., Kurjan, A., Bulet, P., Lemaître, B., Vidal, M., Cordero, J.B. (2019). The antimicrobial peptide defensin cooperates with tumour necrosis factor to drive tumour cell death in Drosophila. Elife. 8, e45061.

Pastor-Pareja, J.C., Wu, M., and Xu, T. (2008). An innate immune response of blood cells to tumors and tissue damage in Drosophila. Dis. Model. Mech. 1, 144–154.

Qin, H., Percival-Smith, A., Li, C., Jia, C.Y., Gloor, G., and Li, S.S. (2004). A novel transmembrane protein recruits numb to the plasma membrane during asymmetric cell division. J. Biol. Chem. 279, 11304–11312.

Remillieux-Leschelle, N., Santamaria, P., and Randsholt, N.B. (2002). Regulation of larval hematopoiesis in Drosophila melanogaster: a role for the multi sex combs gene. Genetics 162, 1259–1274.

Shrestha, R., and Gateff, E. (1982). Ultrastructure and cytochemistry of the cell-types in the tumorous hematopoietic organs and the haemolymph of the mutant lethal (1) malignant blood neoplasm (l(1)mbn) of Drosophila melanogaster. Dev. Growth Differ. 24, 83–98.

Silverman, N., Zhou, R., Stöven, S., Pandey, N., Hultmark, D., and Maniatis, T. (2000). A Drosophila IkappaB kinase complex required for Relish cleavage and antibacterial immunity. Genes Dev. 14, 2461–2471.

Srivastava, A., Pastor-Pareja, J.C., Igaki, T., Pagliarini, R., and Xu, T. (2007). Basement membrane remodeling is essential for Drosophila disc eversion and tumor invasion. Proc. Natl. Acad. Sci. U S A 104, 2721–2726.

Stofanko, M., Kwon, S.Y., and Badenhorst, P. (2008). A misexpression screen to identify regulators of Drosophila larval hemocyte development. Genetics 180, 253–267.

Stöven, S., Ando, I., Kadalayil, L., Engström, Y., and Hultmark, D. (2000). Activation of the Drosophila NF-κB factor Relish by rapid endoproteolytic cleavage. EMBO Rep. 1, 347–352.

Stöven, S., Silverman, N., Junell, A., Hedengren-Olcott, M., Erturk, D., Engström, Y., Maniatis, T., and Hultmark, D. (2003). Caspase-mediated processing of the Drosophila NF-κB factor Relish. Proc. Natl. Acad. Sci. U S A 100, 5991–5996.

Sun, H., Bristow, B.N., Qu, G., and Wasserman, S.A. (2002). A heterotrimeric death domain complex in Toll signaling. Proc. Natl. Acad. Sci. U S A 99, 12871–12876.

Sykiotis, G.P., and Bohmann, D. (2008). Keap1/Nrf2 signaling regulates oxidative stress tolerance and lifespan in Drosophila. Dev. Cell 14, 76–85.

Tzou, P., De Gregorio, E., and Lemaitre, B. (2002). How Drosophila combats microbial infection: a model to study innate immunity and host-pathogen interactions. Curr. Opin. Microbiol. 5, 102–110.

Uhlirova, M., and Bohmann, D. (2006). JNK- and Fos-regulated Mmp1 expression cooperates with Ras to induce invasive tumors in Drosophila. EMBO J. 25, 5294–5304.

Wang, L., Kounatidis, I., and Ligoxygakis, P. (2014). Drosophila as a model to study the role of blood cells in inflammation, innate immunity and cancer. Front Cell Infect Microbiol. 3, 113.

Valanne, S., Wang, J.H., Rämet, M. (2011). The Drosophila Toll signaling pathway. J Immunol. 186, 649–656.

Xie, X., Hu, J., Liu, X., Qin, H., Percival-Smith, A., Rao, Y., and Li, S.S. (2010). NIP/DuoxA is essential for Drosophila embryonic development and regulates oxidative stress response. Int. J. Biol. Sci. 6, 252–267.

